# A quantitative coordinate system for developmental dynamics

**DOI:** 10.64898/2026.06.09.730858

**Authors:** Jiawei Wang, Jinzheng Ren, Cora Moore, Robert N. Kelsh, Katie McDole, Bianca Dumitrascu, John C. Marioni

## Abstract

Quantitative comparison of morphogenesis across individuals remains a fundamental challenge, as developing embryos vary in shape, orientation and developmental tempo. Moreover, real-time three-dimensional imaging generates large, heterogeneous four-dimensional datasets that are difficult to directly align. As a result, developmental variability is typically described qualitatively rather than measured. Here we introduce STERN, a quantitative framework that learns continuous spatiotemporal representations of morphogenesis directly from in vivo 4D imaging data. By embedding embryos into a shared spatiotemporal space, STERN defines a quantitative developmental coordinate system that enables direct comparison of developmental trajectories across individuals without requiring explicit registration or staging. Applied to mouse embryogenesis, STERN reveals that embryos follow conserved developmental trajectories while progressing at distinct temporal rates, providing a quantitative measure of developmental heterochrony. Extending this framework to zebrafish neural crest light-sheet timelapse imaging, we further show that developmental order is preserved across distinct imaging views even with altered anatomical coverage, supporting the generality of the learned representation across vertebrate imaging contexts. Finally, in developing mouse hearts, where morphogenesis proceeds through subtle and continuously evolving structural changes, STERN resolves fine-scale developmental dynamics at minute-scale temporal resolution that are difficult to localize reproducibly using human experts or general-purpose multimodal AI. Together, these results establish a shared quantitative coordinate system for morphogenesis, in which developmental trajectories become directly comparable across individuals and developmental variability becomes a measurable property.

## Introduction

Mammalian embryogenesis unfolds through coordinated cell division, migration and differentiation, generating complex three-dimensional structures through development^1^. Recent advances in *in toto* light-sheet microscopy now enable continuous three-dimensional imaging of mouse embryonic and organ development at cellular resolution, providing an unprecedented view of morphogenesis in space and time^1, 2^. Yet a fundamental challenge remains: how to quantitatively compare morphogenesis across individuals. Developing embryos exhibit substantial inter-individual variability, and continuous three-dimensional time-lapse imaging further complicates direct comparison by revealing morphogenesis as a high-dimensional and continuously changing process. Consequently, developmental variability is typically described qualitatively rather than measured, limiting our ability to extract generalizable principles of morphogenesis.

These challenges arise from the combined effects of data complexity and biological variability. First, temporally adjacent frames can be nearly indistinguishable, whereas distant time points differ substantially, complicating spatial and temporal alignment. Second, embryos at nominally equivalent stages vary in shape, orientation, and developmental tempo, reflecting both technical heterogeneity (e.g., mounting and imaging orientation) and intrinsic inter-individual heterogeneity^3^. Addressing these challenges requires representations that capture global morphological structure while remaining robust to natural variation^3^.

Deep learning approaches have begun to model developmental dynamics in model organisms using primarily two-dimensional image-based representations^4-9^, including inferring developmental time and tempo in zebrafish and *C. elegans* from 2D images^10^. Extending these approaches to in vivo 3D volumetric recordings—particularly in mammals, where imaging orientation, 3D geometry and developmental tempo vary across individuals—remains a fundamental challenge. Solving this problem requires methods capable of learning continuous spatiotemporal representations from volumetric data that enable direct comparison of morphogenesis across individuals.

Here we introduce STERN (Spatio-Temporal Embedding for Real-time morphogeNesis), a 4D morphogenesis embedding framework that learns continuous spatiotemporal structure directly from in vivo 3D recordings. By embedding embryos into a shared spatiotemporal space, STERN defines a quantitative developmental coordinate system that enables direct comparison of morphogenesis across individuals. Technically, STERN combines global visual priors from a pretrained Swin Transformer^11^ with local spatiotemporal constraints learned through contrastive training^12, 13^, yielding embeddings that capture both large-scale morphological organization and fine-grained developmental continuity. By slicing each 3D volume into anatomically ordered 2D images and training on millions of spatially and temporally paired slices, STERN learns a continuous embedding space that reflects the intrinsic structure of mammalian morphogenesis.

Within this coordinate system, morphogenesis can be compared quantitatively across individuals, enabling alignment of full 3D developmental trajectories and measurement of developmental heterochrony—embryos follow similar morphogenetic paths but at distinct temporal rates, consistent with natural variability in mammalian development. Across individuals, aligned trajectories reveal recurrent morphological states, demonstrating that the learned coordinate system captures biologically meaningful developmental features directly from raw imaging data without segmentation. Extending beyond mammalian embryogenesis, we generated a zebrafish neural crest light-sheet timelapse dataset and showed that STERN preserves coherent developmental progression across distinct imaging views despite substantial differences in anatomical coverage and local appearance, suggesting that the learned representation captures developmental organization that is robust across imaging contexts and vertebrate developmental systems. In more challenging settings characterized by subtle and continuously evolving morphology, illustrated in developing mouse hearts, the learned representation enables consistent resolution of fine-scale developmental dynamics beyond what can be reproducibly resolved through expert manual inspection or general-purpose multimodal AI. Together, these results establish a unified framework for quantitative morphogenesis, providing a principled basis for comparing developmental processes across individuals and systems.

## Results

### A continuous 4D representation of mammalian morphogenesis

Our analysis begins with long-term, in toto, light-sheet recordings generated by the Keller laboratory, which include multiple embryos imaged over ∼2.5 days of development^1^. We focused on four representative embryos (A–D) captured from E(Embryonic day)6.0 to E8.5 at five-minute intervals, yielding a continuous 3D record of gastrulation and early organogenesis^1^. For each time point, the volumetric image was sliced into 2D cross-sections along the observation axis (Fig. 1a-b and Methods), producing an anatomically ordered series of images that can, in principle, be compared across individuals.

**Fig. 1.**
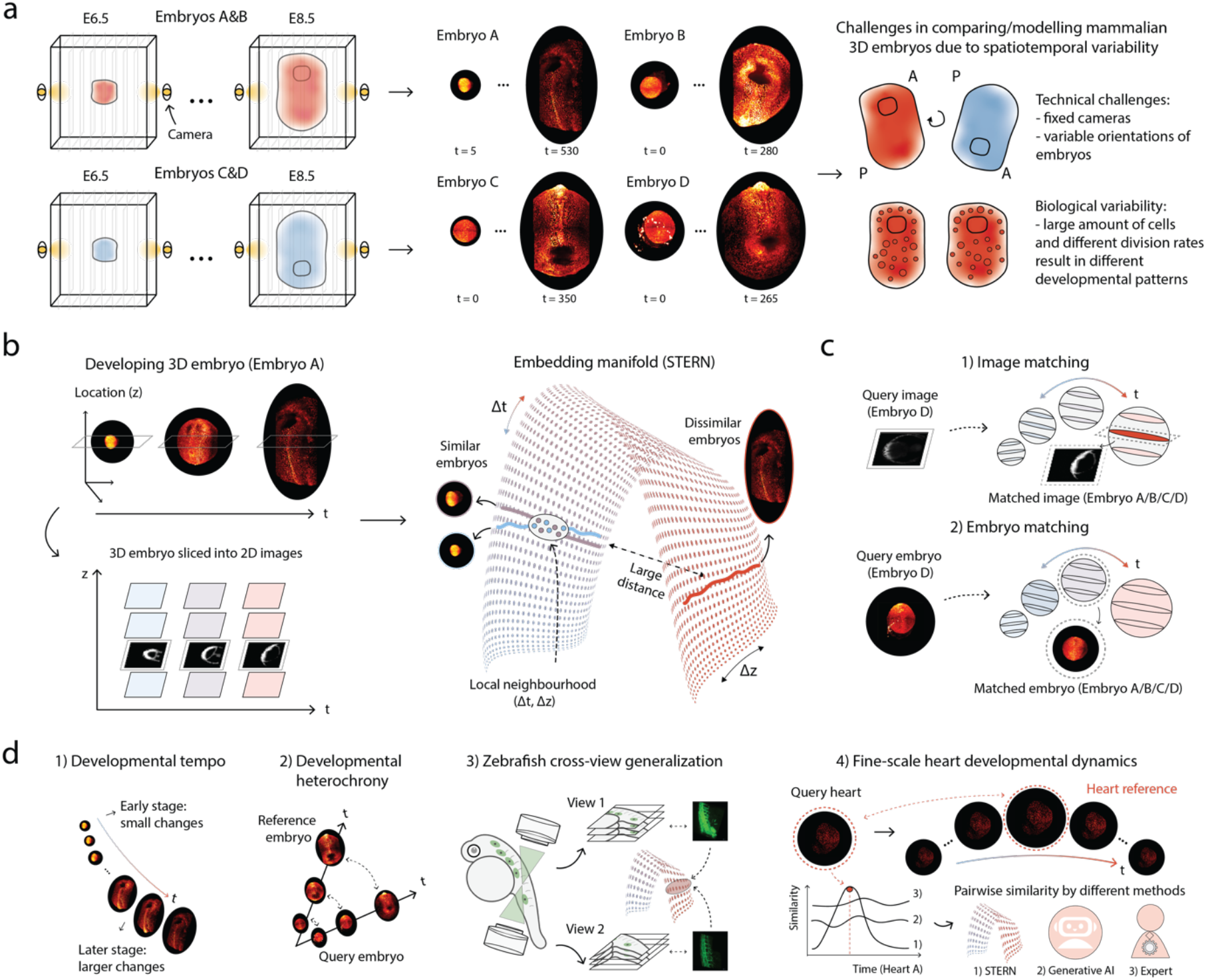
Workflow overview. **a**, Illustration of the process for generating in toto 3D imaging data of mouse embryos (observation direction: Anterior (A) and Posterior (P)), and the challenges encountered in comparing and modelling mammalian 3D embryos due to spatiotemporal variabilities. **b**, The left panel shows a simplified depiction of the in toto 4D imaging data capturing real-time mouse embryo development, illustrating how each 3D volume is sliced into ordered 2D cross-sections along the observation axis (z-location) at every time point. Example slices are shown for illustration. The right panel shows that STERN converts these slices into embeddings that capture inherent spatiotemporal information related to tissue morphogenesis. Embryos that are close in time exhibit similar morphology and therefore map to nearby regions in the embedding manifold, while dissimilar embryos map further apart. **c**, Direct applications of the embeddings include: (1) matching 2D image slices across embryos, and (2) matching 3D embryo volumes by comparing the aggregated similarity of their constituent slices. **d**, Downstream analyses enabled by STERN include quantification of developmental tempo, measurement of developmental heterochrony, cross-view generalization across zebrafish imaging views, and analysis of fine-scale heart developmental dynamics. Images are displayed at different scales throughout Fig. 1 for visual clarity and are not intended for direct size comparison.

A central challenge is that these data contain both global and local sources of variation (Fig. 1a). Globally, embryos differ in overall morphology, imaging orientation, and acquisition conditions. Locally, adjacent time points differ only subtly, yet these fine-scale differences encode the underlying developmental progression. Rather than treating these sources of variability as independent technical obstacles, we sought to learn a representation that organizes them within a shared spatiotemporal structure. Our goal was to learn embeddings that capture this multiscale structure—mapping slices that are similar in space or time to nearby locations in a shared representation space (Fig. 1b).

Classical feature extractors (e.g., SIFT^14^, ORB^15^) and pretrained convolutional or transformer models capture certain structural regularities, but proved insufficient for representing the continuous, multiscale nature of mammalian morphogenesis: classical approaches were overly sensitive to imaging variability, while generic models, trained on discrete object categories, tended to overlook gradual temporal progression. Conversely, contrastive models trained from scratch tended to capture local continuity but failed to recover coherent global organization.

To overcome these limitations, STERN integrates global morphological cues with locally enforced temporal and spatial continuity. The pretrained backbone provides stable representations that are robust to imaging variability, while the contrastive objectives shape a smooth developmental trajectory by encouraging adjacent slices to remain close in the embedding space (Extended Data Fig. 1). To anchor the embedding in biological continuity, we constructed millions of spatially and temporally paired slice examples: slices from nearby z-planes or adjacent time points served as positive pairs, whereas distant slices served as negatives. This dense pairing scheme provides supervisory signals that capture both large-scale tissue organization and short-timescale morphological changes, enabling the model to embed morphogenesis as a continuous trajectory.

Together, these components allow STERN to learn invariant principles of mammalian morphogenesis rather than embryo-specific appearance, while remaining sensitive to subtle morphogenetic changes. Instead of organizing embryos into discrete stages or categories, STERN recovers a continuous developmental manifold in which related morphological states occupy nearby regions of the embedding space. The resulting continuous and generalizable 4D representation was validated across multiple benchmarks and serves as the foundation for all subsequent comparative and quantitative analyses (Fig. 1c-d).

### STERN identifies corresponding developmental states across embryos

#### Recovering developmental correspondence across 2D anatomical sections

To rigorously benchmark the ability of STERN to identify corresponding anatomical sections across independent individuals, we first evaluated 2D slice matching using a held-out biological replicate. After training and fine-tuning models on Embryos A–C (Supplementary Fig. 1a–c), we used Embryo D to define six “query slices” spanning early, mid, and late developmental stages (Fig. 2a). We computed the pairwise distances between these queries and all slices in the reference dataset—using the similarity metric appropriate for each representation (Methods)—and visualized the spatiotemporal distribution of the top 50 matches to assess the global coherence of the mappings (Fig. 2b).

**Fig. 2.**
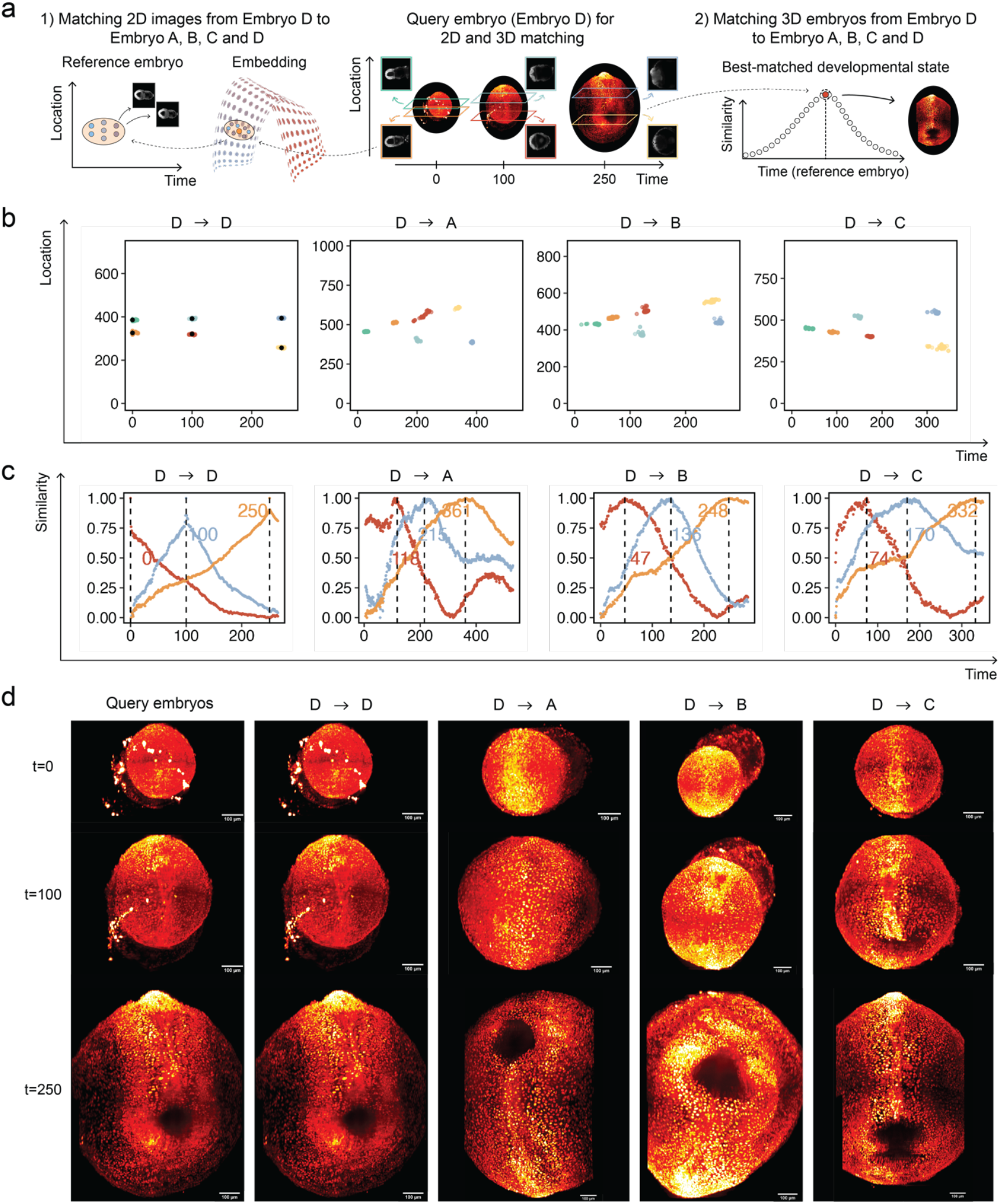
Analysis of 2D image matching using our final model with extension to 3D volumetric alignment. **a**, Schematic of the 2D slice and 3D embryo matching strategy. The middle panel shows three query 3D volumes selected from Embryo D, spanning early, mid, and late time points along the developmental trajectory, which are subsequently used for 3D volumetric matching. From Embryo D, six images are selected as “query images” (query slices). For 2D slice matching, the Euclidean distances between each query slice and all slices in a reference embryo (Embryo A, B, C, or D) are computed, and the top 50 closest matches are visualized as a spatiotemporal scatter plot (left panel). The six query slices are chosen to exhibit clear morphological differences across time. For 3D embryo matching (right panel), the similarity between a query volume and a reference trajectory is computed by aggregating slice-level nearest-neighbour distances into a single global similarity score (Methods). **b**, Matching results for the six distinct query images. “D->D” denotes matching query slices against Embryo D itself, while “D->A/B/C” represents matching query slices against Embryos A, B, and C respectively. This definition applies to all main, extended, and supplementary figures. Each column in the visualization represents the spatial and temporal distributions of the top 50 matching slices for the 6 query slices. Notably, in the “D->D” case, the location of the query slice is denoted by a black dot, serving as an anchor point for reference purposes. **c**, 3D similarity curves for three query time points from Embryo D (t=0, 100, 250) matched against the full developmental trajectories of Embryos A, B, C and D. Vertical dashed lines mark the global maximum (best match), identifying the precise developmental equivalent in the reference embryo. **d**, Visual comparison of the best-matched 3D volumes. Rows represent the three developmental stages (t=0, 100, 250). Columns display the query volume (Embryo D), the self-match (D->D), and the retrieved best matches from Embryos A, B, and C, visualized in an anterior orientation using volumetric rendering.

Across all benchmarks, deep learning approaches consistently outperformed classical feature extractors (SIFT, ORB). This superiority was evident across three metrics: spatiotemporally, deep learning methods produced stable, tight distributions of the top 50 matches compared to the diffuse scatter of classical methods (Extended Data Fig. 2); quantitatively, they achieved significantly higher silhouette widths, indicating better-formed clusters (Extended Data Fig. 3); and qualitatively, the top three retrieved matches consistently exhibited coherent anatomical structures, whereas classical methods frequently returned unrelated slices (Extended Data Fig. 4). Among the deep learning architectures, the Siamese Swin Transformer achieved the highest fidelity, forming six well-separated clusters corresponding to the six query slices (Fig. 2b; Extended Data Fig. 3). Qualitative inspection of the top-retrieved candidates confirmed that the Siamese Swin Transformer consistently identified slices with superior morphological fidelity—closely matching the query images in shape, scale, and structural composition compared to alternative methods.

Beyond model performance, the learned embeddings automatically revealed intrinsic spatiotemporal properties of the dataset. First, the cross-embryo mappings detected global geometric inconsistencies: when matching Embryo D to Embryos A and B, the spatial trajectory of the matches was inverted relative to the query (Fig. 2b; Extended Data Fig. 5). This inversion correctly reflects the physical reality that Embryos A and B were mounted in the opposite anterior-posterior orientation compared to C and D—a biological ground truth recovered purely from the embedding space without prior geometric alignment. Second, the temporal correspondences preserved the global developmental sequence across embryos (Extended Data Fig. 6), while their precision varied by developmental stage. Matches for early gastrulation queries (t=0) were distributed across broader temporal ranges in the reference embryos A–C, whereas matches for mid-to-late stages (t=100, 250) were confined to narrower temporal windows (Fig. 2b; Extended Data Fig. 2), indicating tighter temporal correspondence at later developmental stages. This disparity reflects the changing morphological complexity of the embryo: the early epiblast is a relatively featureless cylinder where adjacent time points are morphologically similar, creating ambiguity. In contrast, later stages involve rapid, distinct structural changes—such as neural fold elevation^16^—that impose strict temporal constraints on anatomical correspondence, allowing STERN to pinpoint developmental stages with high precision.

#### Extending developmental correspondence to full 3D embryo trajectories

While 2D matching validates local feature quality, characterizing the holistic state of an organism requires integrating information across the entire volumetric structure. We therefore extended our evaluation to 3D volumetric matching by integrating information across slices. To quantify similarity between two embryo volumes, we aggregated slice-level nearest-neighbour correspondences across a query volume and a reference volume into a normalized volume-level similarity score that captures overall 3D morphological correspondence. Repeating this comparison across all time points of a reference embryo generated a similarity trajectory whose peak identifies the most morphologically corresponding developmental state for each query volume (Fig. 2a; Methods).

When querying time points from Embryo D against the developmental trajectories of Embryos A, B, and C, the Siamese Swin Transformer generated smooth, unimodal similarity curves with clear peaks, enabling precise identification of developmental equivalents (Fig. 2c; Extended Data Fig. 7). Notably, the model remained robust to non-biological noise; for instance, despite irregular variations in non-peak regions caused by rigid transformations in Embryo A, the model successfully resolved a clear global maximum (Fig. 2c). This indicates that STERN learns a continuous developmental manifold, allowing developmental progression to be represented as a continuous trajectory.

Biological inspection of the top-matched 3D volumes confirmed that STERN effectively tracks complex, non-linear morphogenetic transitions (Fig. 2d). At early stages (t=0), the model correctly mapped the solid, cylinder-like epiblast structure across individuals. At intermediate stages (t=100), it successfully identified the formation of the proamniotic cavity, matching these specific topological changes to their equivalents in the reference embryos. By the later stages (t=250), the model resolved distinct internal cavities and surface peaks associated with the foregut pocket and neural fold elevation (Fig. 2d). Furthermore, the 3D alignment confirmed the orientation differences detected in 2D—aligning the anterior foregut pocket of Embryo D to the spatially inverted equivalents in Embryos A and B. These results demonstrate that STERN integrates distributed morphological cues to resolve fine-grained 4D variability, aligning developmental trajectories even when embryos differ substantially in angle, size, or partial visibility.

### Quantitative alignment of 3D trajectories reveals developmental tempo heterochrony

To quantify the intrinsic pacing of mammalian morphogenesis, we first analysed the dynamics within individual developmental trajectories. By computing the pairwise similarity between all time points for a single embryo, we generated a temporal self-similarity matrix (Fig. 3a-b; Extended Data Fig. 8). Unlike the checkerboard patterns characteristic of discrete developmental stages reported in *C. elegans* or zebrafish, the matrix for mouse embryogenesis reveals a continuous gradient of morphological change. The varying width of the high-similarity diagonal provides a readout of developmental tempo: the broad diagonal at early time points reflects a relatively slow morphogenetic pace during early gastrulation, whereas the narrowing diagonal at later stages indicates a rapid acceleration in morphological restructuring as the embryo undergoes neurulation and early organogenesis (Fig. 3b; Extended Data Fig. 8).

**Fig. 3.**
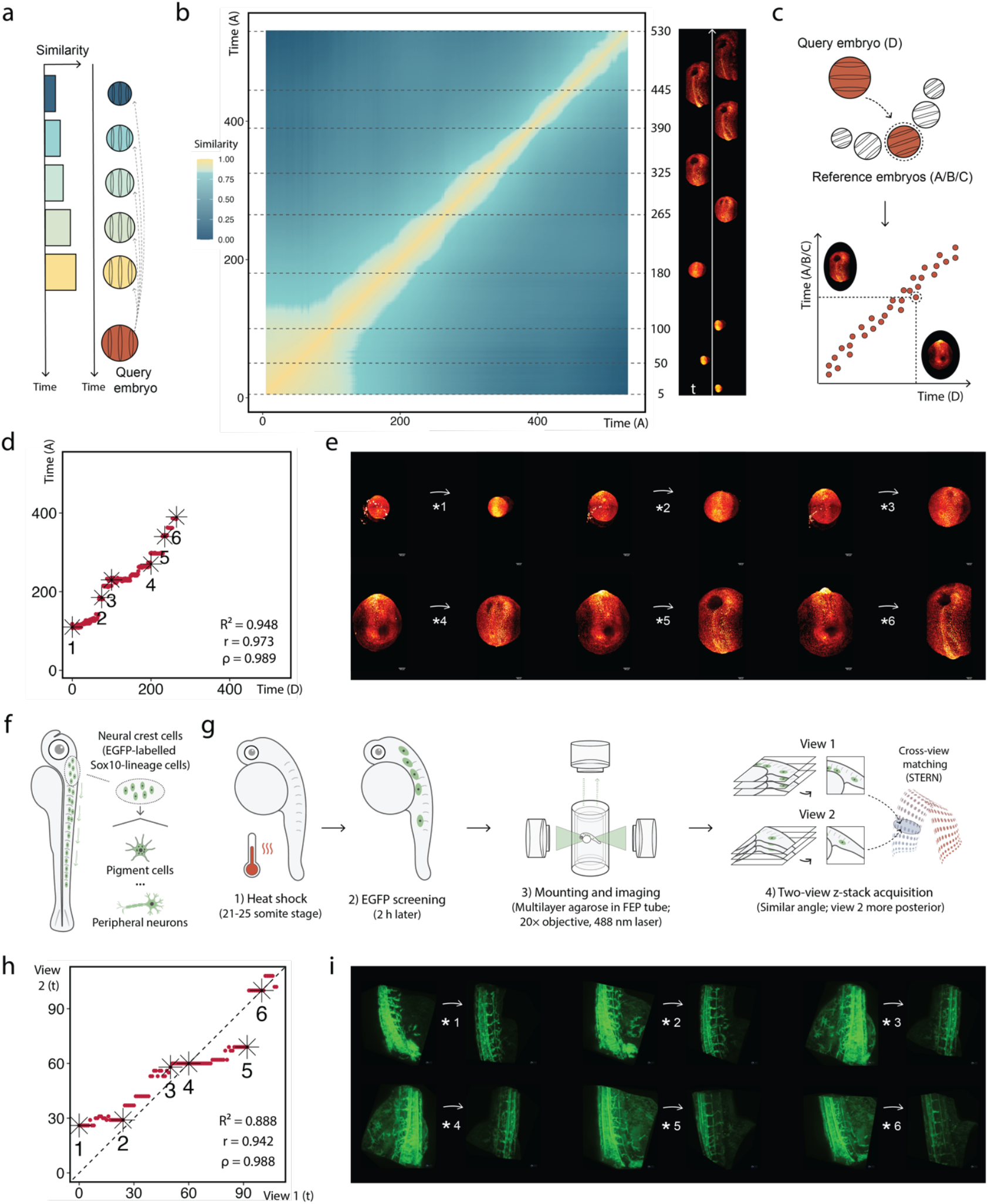
Quantitative alignment of developmental trajectories across embryos and imaging views. **a**, Strategy to generate a similarity matrix. For a query embryo, its similarities against embryos at every time point preceding the query embryo were calculated. These fill half of a similarity matrix, and the rest is symmetrically filled within the similarity matrix. **b**, The similarity matrix for Embryo A as an example, with pairwise comparisons of embryos at every developing time point. High similarity between neighbouring time points delineates clusters corresponding to developmental epochs. Dashed lines mark representative time points during embryo development, with the corresponding embryos shown to track the changes. **c**, Illustration of the cross-embryo temporal alignment strategy. Each 3D volume in the query Embryo D is matched to the closest morphological equivalent in a reference embryo (Embryo A, B, or C). **d**, Comparative analysis of temporal correlation between Embryos D and A. The scatter plot shows the time series of the best-matched volumes, with six representative matched pairs marked by stars (*1–*6). The deviation from linearity indicates heterochrony (R^2^ = 0.948, Pearson r = 0.973, Spearman ρ = 0.989). **e**, Representative matched 3D embryo volumes corresponding to the pairs marked in **d**. For each pair, the query embryo (Embryo D) is shown on the left and the matched reference embryo (Embryo A) on the right. **f**, Schematic overview of the zebrafish neural crest imaging dataset. EGFP-labelled Sox10-lineage neural crest cells were imaged during zebrafish development. **g**, Experimental workflow for zebrafish neural crest light-sheet imaging and cross-view developmental state matching. Two partially overlapping z-stack views were acquired from the same embryo, with the second view positioned more posteriorly along the body axis. Time points from both views were embedded using STERN and compared across the full timelapse to identify corresponding developmental states. **h**, Cross-view temporal alignment between the two zebrafish neural crest imaging views. For each query time point in view 1, all time points in view 2 were evaluated to identify the best-matched developmental state. The scatter plot shows the best-matched time point identified in view 2 for each query time point in view 1. The dashed line indicates the identity relationship. Pearson correlation, Spearman rank correlation and the coefficient of determination from a linear fit are shown. Six representative matched pairs are marked by stars (*1–*6). **i**, Representative cross-view matched developmental states from the six pairs marked in **h**. For each pair, the query state from view 1 is shown on the left and the best-matched state from view 2 is shown on the right. Images are shown as volume-rendered 3D z-stack volumes. Viewing angles were manually adjusted into approximately equivalent viewing perspectives for visualization only and were not used for STERN-based matching. Corresponding maximum-intensity projection views are provided in Extended Data Fig. 10.

We next aligned the full 3D developmental trajectory of embryo D against embryos A, B and C to assess how this pacing is conserved across individuals (Fig. 3c). In all pairwise comparisons, we observed a strong positive relationship between the time points of morphologically matched volumes (Fig. 3d; Extended Data Fig. 9a,c). This indicates that the fundamental sequence of developmental events—from gastrulation through neural fold elevation—is robustly conserved across individuals. Full volumetric alignment proved substantially more robust than local slice-based matching: while 3D matching recovered a clear linear progression, 2D matches frequently showed temporal heterogeneity owing to orientation mismatches and local ambiguities (Extended Data Fig. 5), underscoring the importance of volumetric analysis for resolving true developmental correspondence.

Although the sequence of events was conserved, closer inspection revealed subtle but reproducible heterochrony—variation in the rate at which embryos traverse this sequence. The alignment between embryos D and B served as an approximate isochronous baseline, displaying a highly linear relationship (R^2^ = 0.988; Extended Data Fig. 9a), consistent with both embryos progressing at nearly identical speeds. Relative to this baseline, the alignment of embryo D to embryo A deviated noticeably from linearity (R^2^ = 0.948; Fig. 3d). In the developmental window between waypoints *3 and *4 (Fig. 3d), the curve flattens, suggesting a local delay in the progression of embryo D relative to embryo A through this morphogenetic phase. Visual inspection of the matched 3D volumes at these waypoints supports this interpretation, showing that both embryos pass through comparable morphological states— forming similar internal cavities and surface structures—but over different temporal intervals (Fig. 3e). Together, these results suggest that embryos follow conserved developmental trajectories, while progressing through them at subtly different temporal rates.

Quantitative analysis reinforces the distinction between conserved sequence and flexible tempo. Across all embryo pairs, Spearman correlations remained consistently high (ρ ≈ 0.99), confirming that the rank order of developmental events is preserved (Fig. 3d; Extended Data Fig. 9a,c). In contrast, Pearson correlations and linear fits varied across pairs, with lower values for D→A and D→C than for the D→B baseline, quantifying departures from uniform timing. This combination—highly conserved rank agreement but varying linearity—indicates modest yet consistent tempo heterochrony. Such variability suggests that while the overarching developmental program is stable, its global pace can vary, potentially allowing embryos to buffer stochastic fluctuations in cellular dynamics or environmental conditions without perturbing the critical order of morphogenetic checkpoints.

### Developmental trajectories are conserved across zebrafish imaging views

To test whether the learned developmental coordinate system generalizes beyond mammalian embryo imaging, we generated a zebrafish neural crest light-sheet timelapse dataset in which EGFP-labelled Sox10-lineage cells were imaged during development (Fig. 3f,g; Methods). Unlike the mouse embryo datasets, which capture global embryo-scale morphology, this dataset records a spatially restricted, lineage-labelled migratory cell population within a local posterior trunk/anterior tail field of view (Fig. 3f). This setting therefore introduces a distinct correspondence challenge: developmental progression must be inferred from partial and locally evolving morphology rather than from whole-embryo organization.

Without additional retraining or view-specific optimization, cross-view matching recovered a highly ordered temporal relationship between acquisition time points in the two views, with near-monotonic preservation of developmental progression across the timelapse (Spearman ρ = 0.988; Fig. 3h). Matched pairs near the identity relationship exhibited highly consistent three-dimensional organization of Sox10-positive neural crest populations despite differences in imaging geometry and anatomical coverage between the two views (Fig. 3i). Even modest off-diagonal matches, including *1 and *5, exhibited highly similar spatial organization of Sox10-positive neural crest populations despite occurring away from the identity relationship (Fig. 3i). Comparison with the same-time-point volumes in the second view showed that the identified best matches and the corresponding same-time-point volumes often exhibited highly similar three-dimensional organization, suggesting that local deviations from the identity relationship occur within intervals of gradual morphological change (Extended Data Fig. 10a,b). Together, these results demonstrate that developmental trajectories remain conserved despite partial anatomical overlap and substantial differences in imaging geometry.

### STERN resolves fine-scale cardiac dynamics beyond expert reproducibility

Real-time imaging of early mouse cardiogenesis presents a distinct test of developmental correspondence. Unlike whole-embryo morphogenesis, where large-scale geometric transitions provide strong global cues, early cardiac morphogenesis proceeds through subtle, continuous deformations over minute-scale intervals. Individual 2D projections and cross-sections can therefore remain visually similar across adjacent or nearby frames, making the precise localization of developmental state challenging for both manual inspection and automated image comparison (3-min sampling; Fig. 4a).

**Fig. 4.**
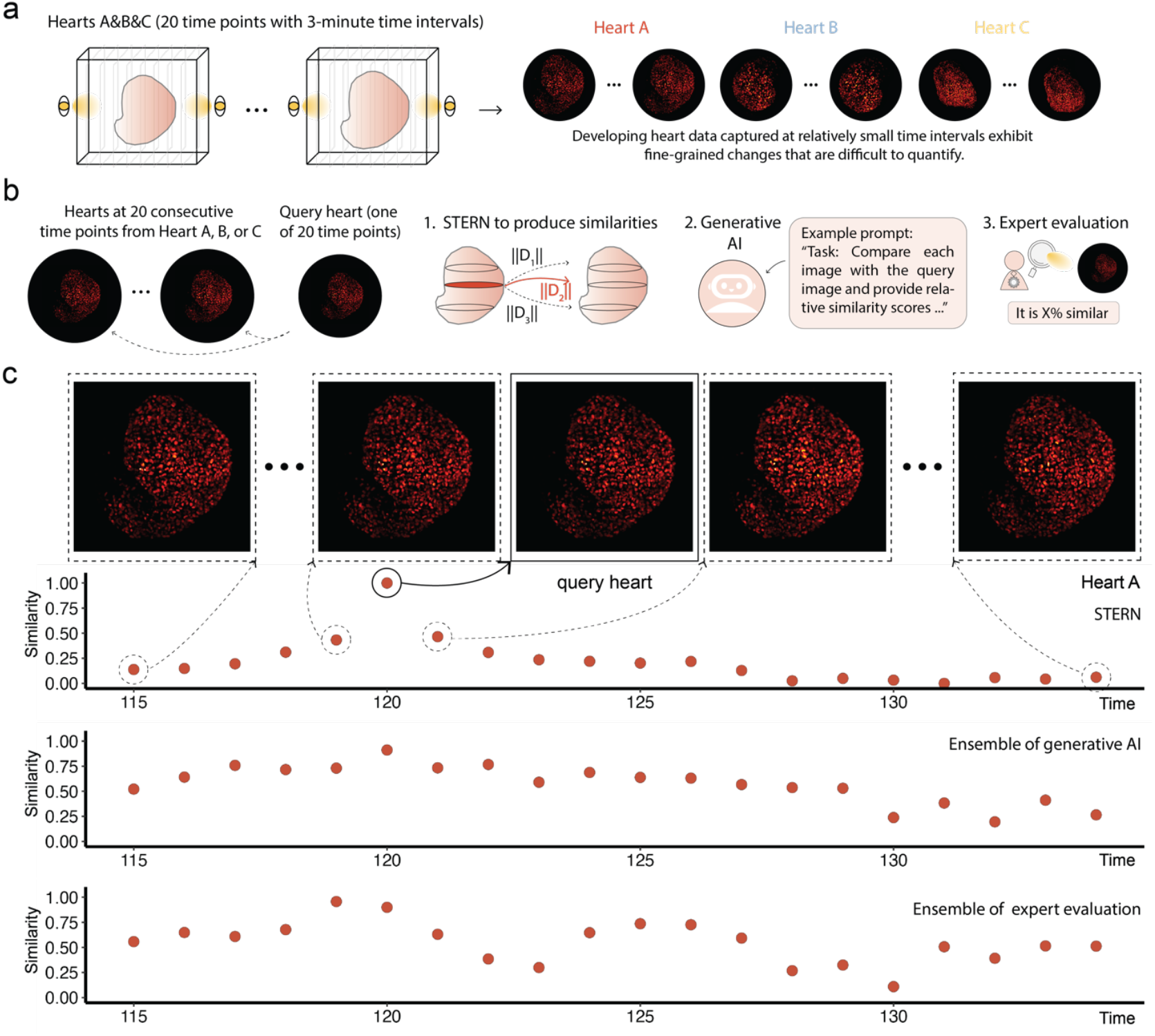
Volumetric alignment of fine-scale cardiac morphogenesis. **a**, Data acquisition for fine-scale organ dynamics. Three independent developing mouse hearts (A, B and C) were imaged at 3-min intervals over 20 time points. Adjacent frames exhibit subtle morphological changes, creating a challenge for precise temporal localization. **b**, Benchmark strategy for resolving cardiac developmental state. A query heart volume was matched against the full temporal sequence using three approaches: STERN volumetric embedding, general-purpose multimodal AI evaluation and blinded expert evaluation. **c**, Comparison of temporal similarity trajectories for Heart A. The three lower panels show normalized similarity trajectories produced by STERN (top), the multimodal AI ensemble (middle), and the expert ensemble (bottom), respectively, when matching a query heart volume at time point t = 120 against all time points in Heart A. Higher scores indicate greater similarity to the query. The image panel above shows representative reference volumes sampled from distinct regions of the STERN trajectory, including highly similar and weakly similar matches relative to the query heart volume. Dashed lines connect selected trajectory positions to their corresponding representative images. All scores were normalized to range from 0 to 1.

We first examined whether STERN could recover developmental correspondence from individual 2D sections of three independent hearts (A–C), without heart-specific retraining. Querying a single slice against its developmental trajectory produced broad, plateau-like similarity profiles rather than sharply localized peaks (Extended Data Fig. 11a). This pattern reflects the biological continuity of cardiac morphogenesis at minute resolution: adjacent frames often represent genuinely similar developmental states. Consequently, although slice-based comparisons reliably recovered anatomically corresponding regions across datasets, they provided limited temporal precision for resolving closely spaced cardiac states (Extended Data Fig. 11a-b).

We therefore extended the analysis from local sections to full 3D heart volumes (Fig. 4b). Because early cardiac morphogenesis unfolds through weak but coordinated deformations distributed across the tissue, volumetric integration provides a stronger readout of developmental progression than any single view or section. When full 3D volumes were embedded using STERN, similarity–time trajectories became smooth and locally peaked, with each query volume showing a clearly defined maximum at the corresponding developmental time point (Fig. 4c, first row; Extended Data Fig. 12). Thus, minute-scale developmental order can be recovered when distributed morphological cues are integrated across the whole organ.

To place this temporal resolution in context, we compared STERN with blinded evaluations from general-purpose multimodal AI systems and domain experts (Fig. 4b; Methods). The AI baseline included GPT-5.5 (standard thinking), Claude Sonnet 4.6 and Gemini 3 Flash, each tasked with scoring visual similarity between a query heart image and anonymized reference frames. Individual AI models showed variable behaviour across heart datasets (Supplementary Fig. 2); although GPT-5.5 standard thinking recovered relatively coherent temporal structure in Hearts A and B, this was not reproduced consistently across all datasets, particularly in Heart C. We therefore used an ensemble AI trajectory in the main figure to summarize the shared tendency across models without over-interpreting model-specific behaviour (Fig. 4c, middle row; Extended Data Fig. 12). The ensemble captured broad visual similarity around the corresponding developmental window but produced less smooth and less consistently localized trajectories than STERN. Human experts similarly identified the approximate temporal neighbourhood of the query, but consensus trajectories were less sharply localized and individual assessments varied across evaluators and datasets (Fig. 4c, bottom row; Supplementary Fig. 3; Extended Data Fig. 12).

Together, these results show that STERN resolves fine-scale cardiac developmental states more reproducibly than either general-purpose multimodal AI or expert visual assessment. STERN integrates weak, distributed morphological cues across 3D volumes to recover temporally coherent developmental trajectories. This extends the coordinate-system framework from whole-embryo morphogenesis to organ-level dynamics, demonstrating that STERN can quantify developmental progression in regimes dominated by subtle, continuously evolving structural change.

## Discussion

Understanding how complex organisms emerge from coordinated cellular behaviours requires frameworks that can compare development quantitatively across individuals despite variability in shape, orientation and developmental tempo. Here, we show that morphogenesis can be embedded within a shared quantitative coordinate system learned directly from real-time volumetric imaging data. This representation enables developmental trajectories to be aligned without explicit staging or registration, transforming qualitative descriptions of variability into measurable, comparable quantities.

Across mouse embryogenesis, this coordinate system reveals a reproducible pattern of mammalian development: while the sequence of morphogenetic events is highly conserved, the rate at which embryos progress through this sequence varies across individuals. This structured heterochrony— characterized by highly conserved developmental order alongside subtle but measurable variation in tempo—suggests that morphogenesis follows a robust developmental programme that accommodates flexibility in timing. Such variability is difficult to resolve using conventional staging frameworks or manual inspection, which typically rely on a limited set of morphological criteria and often obscure continuous developmental dynamics. By contrast, continuous representations capture both global progression and local transitions, enabling developmental tempo to be quantified as an intrinsic property rather than inferred post hoc.

The cross-view zebrafish neural crest analysis further demonstrates that this coordinate system extends beyond globally organized embryo morphology to settings involving partial views with differing anatomical coverage. The preservation of developmental order across such views suggests that the learned representation captures intrinsic aspects of developmental progression that are robust to changes in imaging geometry and local appearance. Together, these results support the view that morphogenesis, despite its apparent complexity, can be organized within a coherent and comparable structure across vertebrate developmental systems.

STERN also generalizes to developmental regimes characterized by subtle and continuously evolving morphology. In developing mouse hearts, where morphogenesis is dominated by subtle, continuous deformations at minute-scale resolution, STERN remains sensitive to subtle but coordinated developmental progression that is difficult to localize reproducibly through manual inspection or general-purpose multimodal AI. This highlights the advantages of learning directly from spatiotemporally structured data, enabling representations that capture distributed, dynamically evolving patterns underlying developmental state.

More broadly, our results suggest that morphological dynamics may serve as a quantitative readout of underlying developmental state. While STERN operates purely on imaging data, the consistency of the recovered trajectories and their alignment across individuals imply that spatial structure encodes rich information about developmental progression. This raises the possibility of linking morphological and molecular descriptions of development within a unified framework. Recent advances in spatial transcriptomics and multimodal single-cell profiling provide complementary snapshots of gene expression^17-19^, but typically lack continuous temporal resolution. Integrating such data with spatiotemporal embeddings could enable direct mapping between gene expression dynamics and morphological change, offering a more complete view of developmental processes.

In summary, by embedding morphogenesis within a shared spatiotemporal coordinate system, STERN provides a quantitative framework for comparing development across individuals. Rather than viewing developmental variability solely as a source of analytical complexity, this framework enables variability to be measured, interpreted and potentially linked to underlying biological mechanisms. This perspective opens new avenues for studying the principles that govern developmental dynamics, moving towards a more quantitative and integrative understanding of developmental organization across scales.

## Methods

### Data acquisition

All four embryos were imaged during developmental stages E6.0 days post conception (d.p.c.) to E8.5 d.p.c. every 5 min with a z-step size of 2.031 *μm*, where the image data sets of mouse development, captured using a light-sheet microscope, are stored in the KLB 2.0^1^ lossless compression format. These datasets can be accessed at the Image Data Resource website (https://idr.openmicroscopy.org/webclient/?show=project-502). To facilitate further analysis, we employed a data pre-processing pipeline on the KLB data sets. For each time point, we divided the KLB data into a series of images in the standard JPG format. These images represent cross sections of the embryo and are commonly referred to as z-slices or image slices. Consequently, Embryo A consists of 532 time points, each containing 988 z-slices. Similarly, Embryo B comprises 286 time points with 947 z-slices, Embryo C includes 351 time points with 934 z-slices, and Embryo D encompasses 266 time points with 719 z-slices. By transforming the KLB data into individual image slices in the JPG format, we created a more accessible and manageable dataset for subsequent analysis and processing.

### Dataset construction

The construction of the dataset involved selecting positive and negative samples from four embryos. During the selection of positive and negative samples, addressing the abundance of empty image slices within the data is crucial. Due to the nature of the imaging technology utilized, not all z-slices contain useful information regarding the development of the mouse at each time point. Consequently, including these empty slices in our dataset would be unnecessary for constructing the dataset (Supplementary Fig. 4a). To address the issue of abundant empty image slices in our dataset, we developed a method for establishing the upper and lower z-slice bounds at each time point (Supplementary Fig. 4b). These bounds, denoted as 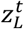 (*z* value of the lower bound) and 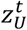 (*z* value of the upper bound), encapsulate the z-slices that predominantly contain relevant embryo content at time point *t*, thereby excluding the empty slices that lie outside these bounds. Manually determining these bounds for each time point would be impractical, while fully automating the process would be overly complex. Instead, we adopted a semi-automated approach by leveraging the temporal stability of the images, where all time points were registered to a common reference frame^1^. The temporal stability allows us to assume there are no significant discontinuities between the bounds from one time point to the next. We started by manually identifying the lower and upper bounds at a few key time points by visually inspecting the images and selecting slices with minimal embryo content, typically located at the embryo’s boundaries. Assume the lower and upper bounds were manually selected at two time points, denoted as, 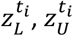 and 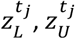. For time points between *t*_*i*_ and *t*_*j*_, we applied linear interpolation to infer the bounds. For any time point *t*_*k*_ between these manually determined points, the lower and upper bounds are calculated as follows:

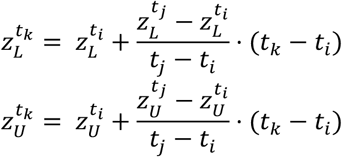

By adopting this approach, we ensured that our dataset exclusively contains z-slices with relevant embryo content, thereby improving the efficiency of the model by excluding empty slices. (Supplementary Fig. 4c)

After defining the upper and lower bounds for each time point, the z-slices were divided into four sections (each containing the same number of z-slices), ordered by increasing z-values, to extract specific embryo content. Sections 2 and 3, located near the centre, primarily captured the embryo’s central content, while Sections 1 and 4, near the lower and upper bounds, mostly contained peripheral content (Supplementary Fig. 5). By organizing the z-slices into these sections, we observed that images in adjacent sections were generally similar, providing a basis for selecting positive and negative samples along the z-axis. Next, we considered the temporal relationship between the slices. We observed minimal differences between corresponding z-slices across consecutive time points. It often took several time points before a noticeable change in shape occurred. This temporal continuity was taken into account during the sample selection process as it helped to refine the selection of positive and negative samples along the temporal axis by considering the temporal relationship between z-slices.

By considering both the proximity of adjacent sections and the temporal continuity, we were able to construct the positive and negative samples. The process began by iterating through all non-empty images across all available time points. For an image at each iteration, referred to as the “original slice,” the remaining images were divided into three classes: a positive class, an ambiguous class, and a negative class. Slices in the positive class were considered candidate positive sample pairings with the original slice, while slices in the negative class were considered candidates for negative pairings. The ambiguous class contained images that were challenging to classify as either similar or dissimilar and could introduce errors if misclassified. By refining these three classes, we ensured the construction of high-quality positive and negative samples. A detailed explanation of the sample selection process is provided in the **Supplementary sections** “***selection of positive samples”*** and “***selection of negative samples”***.

Following the above procedure, we were able to generate positive and negative samples for each of our four embryos. Approximately 1,100,000 samples (including both positive and negative samples) were generated from Embryo A, B, and C. We divided the data into training, validation, and testing sets in a 7:1:2 ratio. Around 130,000 samples generated from Embryo D were used as additional test data and were not including during training and validation. The goal was to demonstrate that our model can effectively handle the inherent variability across experiments.

### Network training details

The Siamese Swin Transformer was designed to extract spatiotemporal information associated with tissue morphogenesis from image slices obtained from developing mouse embryos. The network architecture comprises the following four major components:

1. A data augmentation module was employed to transform any given image *x* randomly, resulting in one related view of the same image, denoted as *x*’. These augmentations were designed to generate diverse and challenging image variations while maintaining essential features for model training.
2. A neural network backbone *f*(·) to extract representation vectors from augmented images. Our framework supports various network architectures; in this case, we used the Swin Transformer^11^, pretrained on the ImageNet-22K dataset^20, 21^. The network produces the representation *h* = *f*(*x*′) = *SwinTransformer*(*x*′), where *h* ∈ *R*^*d*^ is the output from the average pooling layer.
3. A small neural network projection head *g*(·) to map the representations into a space where contrastive loss is applied. Specifically, we employed a 3-layer Multi-Layer Perceptron (MLP) to obtain *z* = *g*(*h*), where *z* ∈ *R*^*k*^.
4. A contrastive loss function^22^ was defined to learn the similarities of images. Given an image pair (*x*_1_, *x*_2_) that is either a positive or negative sample, each image was passed to our backbone and projection head to generate its respective output (*z*_1_, *z*_2_). If *D*_*w*_(*x*_1_, *x*_2_) = ‖*z*_1_ − *z*_2_‖_2_ represents the Euclidean distance between *z*_1_ and *z*_2_, the contrastive loss function is defined as:

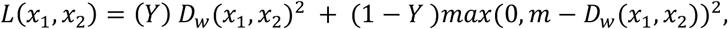

where *Y* is a binary label, with 1 indicating that the input image pair represents a positive sample (the input images are similar), and 0 a negative sample (the input pairs are dissimilar). *D*_*w*_ denotes the Euclidean distance between the embeddings of the two images and *m* is a non-negative parameter that defines a margin. For dissimilar pairs, the loss contribution is only considered if their distance falls within this margin. The goal of the contrastive loss function is to minimize the distance between positive samples while penalizing negative samples whose distance falls within the margin, thereby encouraging separation beyond the margin. By optimizing this loss function, the Siamese Swin Transformer aims to effectively learn features that capture the similarity or dissimilarity between image slices in the context of tissue morphogenesis. After the training is completed, we discard the projection head and solely rely on the backbone to extract features for downstream tasks.

In this work, the Siamese Swin Transformer was trained on the European Bioinformatics Institute (EBI) HPC equipped with an Intel(R) Xeon(R) Gold 6336Y CPU at 2.40GHz and a Tesla V100-PCIE-32GB GPU, using Python (3.7.12) and Pytorch (1.8.0). The training settings for the Siamese Swin Transformer largely aligned with the self-supervised learning approach employed in the study on Swin Transformer^23^. By default, we used the AdamW^24^ optimizer for 50 epochs, with a cosine decay learning rate and 5 epochs of linear warm-up. A batch size of 32, an initial learning rate of 6.25e-07, and a weight decay of 0.001 were employed.

In our data augmentation procedure, two distinct transformations were applied to each image pair input during learning, following the approach outlined in BYOL^25^. These transformations were designed to generate two different views of the same image. For the first transformation, we randomly cropped and resized the image to 224×224 pixels, ensuring that the crop covered between 70% to 100% of the original area. We then applied a series of augmentations, including color jitter (with brightness, contrast, saturation, and hue adjustments of 0.4, 0.4, 0.2, and 0.1, respectively), random grayscale conversion (with a probability of 0.2), Gaussian blur (applied with a sigma range of 0.1 to 2.0 and a probability of 1.0), and horizontal flipping. The image was then normalized using standard ImageNet statistics (mean: [0.485, 0.456, 0.406], std: [0.229, 0.224, 0.225]). The second transformation followed a similar procedure, starting with random cropping and resizing to 224×224 pixels and color jitter with the same parameters. However, the Gaussian blur was applied with a reduced probability of 0.1, and we added a solarization transformation with a probability of 0.2. Horizontal flipping and normalization using ImageNet statistics were also applied. These two augmentations were used to generate the paired views of the images during learning, ensuring that each image in the pair underwent a unique augmentation process. This augmentation strategy was consistent across both training and validation phases.

During training, we used a pretrained Swin Transformer called ‘swin_large_patch4_window7_224_in22k’, obtained from the PyTorch Image Models (timm^26^) library. The default input image resolution was set to 224 x 224. To adapt the pretrained model for our task, we removed the last prediction layer and used the output from the average pooling layer. The output of the backbone is a 1536-dimensional vector, which is subsequently fed into the projection head. The projection head is a 3-layer MLP, where the hidden layers have a dimensionality of 4096 and included the ReLU activation function^27^. The output layer of the MLP is 256-dimensional and does not incorporate the ReLU activation. All layers in the MLP are accompanied by Batch Normalization^28^, following the approach of SimCLR^29^. Regarding the loss function, the margin value (*m*) is set to 1.25. The training details for the Siamese Swin Transformer are provided above. The second model, Siamese Swin Transformer (from scratch), which shares the same architecture, was trained under identical settings. The key difference lies in initializing the Swin Transformer with random parameters instead of using the pretrained weights.

Supplementary Fig. 1a displays the training and validation loss curves for both models. The figure illustrates the convergence behaviour and overall performance of each model throughout the training process. We evaluated both models, along with ResNet and Swin Transformer, on two test sets: 10% of the data from Embryos A, B, and C (denoted as test-ABC) and all samples from Embryo D (denoted as test-D). Supplementary Fig. 1b-c shows the distribution of distances computed between image pairs in the positive and negative samples (see “**2D Image Matching**” section for details on computing distances between images using deep learning methods). The figure demonstrates consistent separation between positive and negative pairs across different test sets. This clear separation between positive and negative sample distributions indicates successful contrastive learning, with positive pairs consistently showing smaller distances compared to negative pairs across all model variants.

### 2D image matching

We benchmarked our Siamese Swin Transformer against the following classical methods and deep learning methods in the 2D image matching. The benchmark is based on similarity measurement between two given images.

For classical methods, we used SIFT and ORB from the OpenCV library^30^ to detect keypoints and compute descriptors for each image. We then compare the keypoints and descriptors of two images using a brute-force matcher (from OpenCV). This matcher pairs the descriptors from the two images and returns the closest matches based on the distances. Depending on the method used (e.g., SIFT, ORB), different distance metrics are applied: for ORB, the Hamming distance is used, while for SIFT, the Euclidean distance is the default. After matching, we apply a ratio test to filter out poor matches. This step ensures that only matches where the distance to the best match is significantly lower than the second-best match are retained. A ratio of 0.75 is used, meaning the distance to the best match must be at least 25% smaller than the next closest match. Once the good matches are filtered, we compute the final similarity score by dividing the number of correct matches (good points) by the total number of matches. This ratio is returned as a similarity score between 0 and 1, where 1 indicates high similarity and 0 indicates low similarity.

For deep learning methods, we examined four models. ResNet-50 pretrained on a pre-processed ImageNet-21K dataset^21^, here, the model takes an image and outputs a feature vector of dimension 2048. A Swin Transformer model (‘swin-large-patch4-window7-224-in22k’) pre-trained on the ImageNet-21k dataset. A plain Swin Transformer model (same architecture as above) trained on our dataset of image pairs, namely Siamese Swin Transformer (from scratch). Finally, a pre-trained Swin Transformer model fine-tuned on our dataset of image pairs, namely Siamese Swin Transformer. The above three Swin Transfomer based models take an image and output a feature vector of dimension 1536. All the features of the four methods are taken as the output from the average pooling layer. As a result, we extracted features for each of the two given images and compute the Euclidean distance as the associated similarity score for 2D image matching.

### Silhouette width

The Silhouette width assesses whether a clustering approach minimizes dissimilarity within clusters and maximizes dissimilarity between clusters^31^. Assuming there exists a clustering into multiple clusters, the Silhouette width for each sample, denoted as *i*, is defined as follows.

1. Calculate *a*(*i*): The average dissimilarity between sample *i* and all other data points in its cluster *A*.
2. For each cluster *C* ≠ *A*, compute *d*(*i, C*): The average dissimilarity between sample *i* and all data points in cluster *C*.
3. Identify cluster *B* with the minimal dissimilarity *d*(*i, B*) to sample *i* : *b*(*i*): = *min*_*C*_*d*(*i, C*). Cluster *B* represents the ‘neighbouring’ cluster to sample *i*.
4. Calculate the Silhouette width s(i) as the scaled difference between the average dissimilarity within the cluster and the average dissimilarity to the neighbouring cluster: s(i) = (b(i) - a(i)) / max(a(i), b(i)).

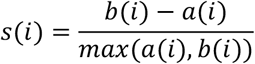

In this study, we used the calculation method from the bluster R package, which approximates the average dissimilarity using the root-mean-squared distance.

### 3D embryo distance computation

To compute the similarity between two mouse embryos, denoted as *E*_*query*_ and *E*_*ref*_, let 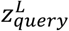 and 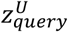 represent the lower and upper bounds of *E*_*query*_, which contain the embryo content. Similarly, let 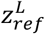 and 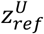 denote the bounds for *E*_*ref*_. We then define the sets of valid slices associated with the query and reference embryos as follows:

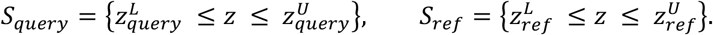

For each slice in both embryos, we extract feature vectors that represent the content of the corresponding slice. Let *f*(*slice*_*z*_) ∈ *R*^*d*^ denote the feature vector for *slice*_*z*_, where *d* is the dimensionality of the feature space.

To compute the similarity between the two embryos, we match slices from the query embryo to their nearest neighbors in the reference embryo based on the Euclidean distance between their feature vectors. The nearest neighbour for a slice *slice*_*query*_ ∈ *S*_*query*_ is the slice *slice*_*ref*_ in the reference embryo that minimizes the Euclidean distance:

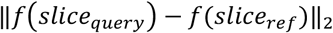

For each *slice*_*query*_ in the query embryo, we compute the minimum distance to the corresponding slice in the reference embryo. The total similarity between the two embryos is defined as the sum of these minimum distances:

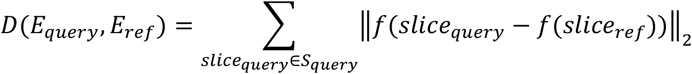

This accumulated distance quantifies the similarity between the two embryos, with a lower value indicating greater similarity.

### 3D embryo matching

For each of the three query embryos from Embryo D, taken at time points 0, 100, and 250 (denoted as *E*_*query*_), we compute the distance between the single query embryo and all embryos within one of the reference sets: Embryo A, B, or C. This results in a set of distances, *D*_*ref*_, where *D*_*ref*_ represents the distances between *E*_*query*_ and all embryos in either Embryo A, B, or C. To convert these raw distances into a more interpretable similarity score ranging from 0 to 1—where a value closer to 1 indicates higher similarity and a value closer to 0 indicates lower similarity—we apply the following normalization for each element in *D*_*ref*_:

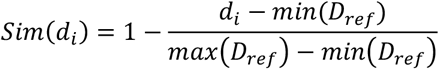

### Ethics statement and zebrafish husbandry

This study was performed with approval from the Ethics Committee of the University of Bath and in accordance with the Animals Scientific Procedures Act (ASPA) 1986 under the Home Office Project Licence P87C67227. Adult zebrafish housed within the University of Bath Fish Facility on a 14 h:10 h light:dark cycle were used to generate embryos through natural matings. Embryos were staged according to Kimmel et al^32^.

### Zebrafish neural crest light-sheet imaging

Zebrafish neural crest timelapse imaging was performed using Tg(Sox10:Cre); Tg(hsp70l:LOXP-DsRed-LOXP-Lyn-EGFP) transgenic embryos on an AB wild-type background. Sox10-lineage cells were labelled through heat-shock-induced EGFP expression. Embryos were heat-shocked at the 21–25 somite stage to induce transgene expression and screened 2 h later for EGFP-positive signal. The EGFP-positive embryo used for this two-view analysis (see below) was mounted in 0.1% agarose containing MS-222 anaesthetic, surrounded by an outer cylinder of 3% agarose to increase confinement within the FEP tube, following a multilayer agarose mounting strategy^33^. Timelapses were acquired every 10 min using a 20× objective and 488 nm laser excitation. For each time point, z-stacks spanning the approximate embryo depth within the field of view were acquired along the dorsal–ventral direction at 1 μm increments. Left and right light-sheet illumination projections were fused using maximum projection using the Zeiss ZEN 3.1 black edition. The cross-view analysis used two related views of the same zebrafish embryo. The two views were acquired from similar imaging orientations, but the second field of view was positioned more posteriorly along the body axis. The analysed two-view dataset captured development from approximately 25 to 44 hours post fertilization (hpf) and contained 109 time points in each view. View 1 comprised 311 z-planes per time point and view 2 comprised 363 z-planes per time point.

### Zebrafish cross-view developmental correspondence analysis

For zebrafish cross-view developmental correspondence analysis, each time point from view 1 was compared against all time points from view 2 using the same STERN-based 3D matching procedure described above. Unlike the mouse embryo analysis, where valid z-slice bounds were defined to restrict the comparison to embryo-containing regions, the zebrafish analysis used the full acquired z-stack for both views. Thus, each comparison used all 311 z-planes from the query time point in view 1 and all 363 z-planes from each time point in view 2. For each query, the time point in view 2 with the lowest accumulated STERN feature distance was selected as the best match.

### Developing mouse heart data and pre-processing

Data from three mouse hearts were recorded at the following developmental stages: Heart A (E8.5–E10.0), Heart B (E8.25–E9.5), and Heart C (E8.75–E9.5). Each heart was imaged during its respective developmental stages every 3 minutes with a z-step size of 2.5 μm, and the resulting datasets, captured using a light-sheet microscope, are stored in .tif format. Due to data size constraints, only 20 time points of imaging data were provided for each developing heart in the public dataset (https://figshare.com/projects/Long-term_live_imaging_of_mouse_embryonic_heart/74532), and these were used in our analysis. The .tif imaging data were converted into a series of images in JPG format. Specifically, Heart A includes time points 115 to 134, with each time point containing 130 z-slices; Heart B includes time points 251 to 270, with each containing 160 z-slices; and Heart C includes time points 101 to 120, each also containing 160 z-slices. We manually determined the lower and upper bounds for each heart dataset due to the small number of time points.

### Dimension reduction on mouse hearts data

We collected all z-slices between the lower and upper bounds for Hearts A, B, and C. Features were extracted from each z-slice using the Siamese Swin Transformer. To reduce the dimensionality of the data, we applied Principal Component Analysis (PCA^34^) followed by t-distributed Stochastic Neighbor Embedding (t-SNE^35^, both from the scikit-learn library^36^). PCA was performed to reduce the data to 50 principal components, capturing most of the dataset’s variance. Following PCA, t-SNE was applied to embed the data into two dimensions. Configured with a perplexity of 40 and 300 iterations, t-SNE transformed the principal components into a two-dimensional space for visualization (Extended Data Fig. 11b). In addition, we analyzed three dimensionality reduction techniques: PCA, t-SNE, and UMAP^37^ (Uniform Manifold Approximation and Projection). Supplementary Fig. 6 presents the results of dimension reduction on Hearts A, B, and C using these methods. PCA reduced the data directly to 2 dimensions (Supplementary Fig. 6, leftmost panel). t-SNE was applied with a perplexity of 40 and 300 iterations to reduce the data directly to 2 dimensions (Supplementary Fig. 6, middle panel). UMAP reduced the data to 2 dimensions using parameters ‘*n_neighbors=40*’ and ‘*min_dist=0*.*6*’, implemented with the umap-learn library (Supplementary Fig. 6, rightmost panel).

### Blinded AI and expert similarity evaluation

To benchmark STERN against human perception and general-purpose multimodal models, we conducted a blinded similarity evaluation using maximum-intensity projections derived from developing 3D heart volumes. For each heart dataset (A–C), one query image was compared against 20 reference images from the same dataset. Reference images were randomized and anonymized prior to evaluation, and no temporal information was provided.

For the AI baseline, three multimodal models (GPT-5.5 standard thinking, Claude Sonnet 4.6 and Gemini 3 Flash) were evaluated using the same image sets. Images were uploaded in three sequential batches consisting of one query image followed by two groups of ten reference images. Models were instructed to assign a similarity score from 1 to 100 to each reference image relative to the query image and to return only the image filename and score (**Supplementary Note 1**). For the expert baseline, five domain experts independently evaluated the same image sets. Experts assigned similarity scores from 1 to 100 to each reference image relative to the query image using standardized spreadsheets. Expert backgrounds included developmental biology, bioimage analysis and related fields; this information was collected to interpret variability across evaluations and is reported in anonymized form (**Supplementary Table S1**). For each dataset, scores were normalized independently within each evaluator using min–max normalization. Consensus AI and expert trajectories were computed by averaging normalized scores across models or experts, respectively.

**Extended Data Fig. 1.**
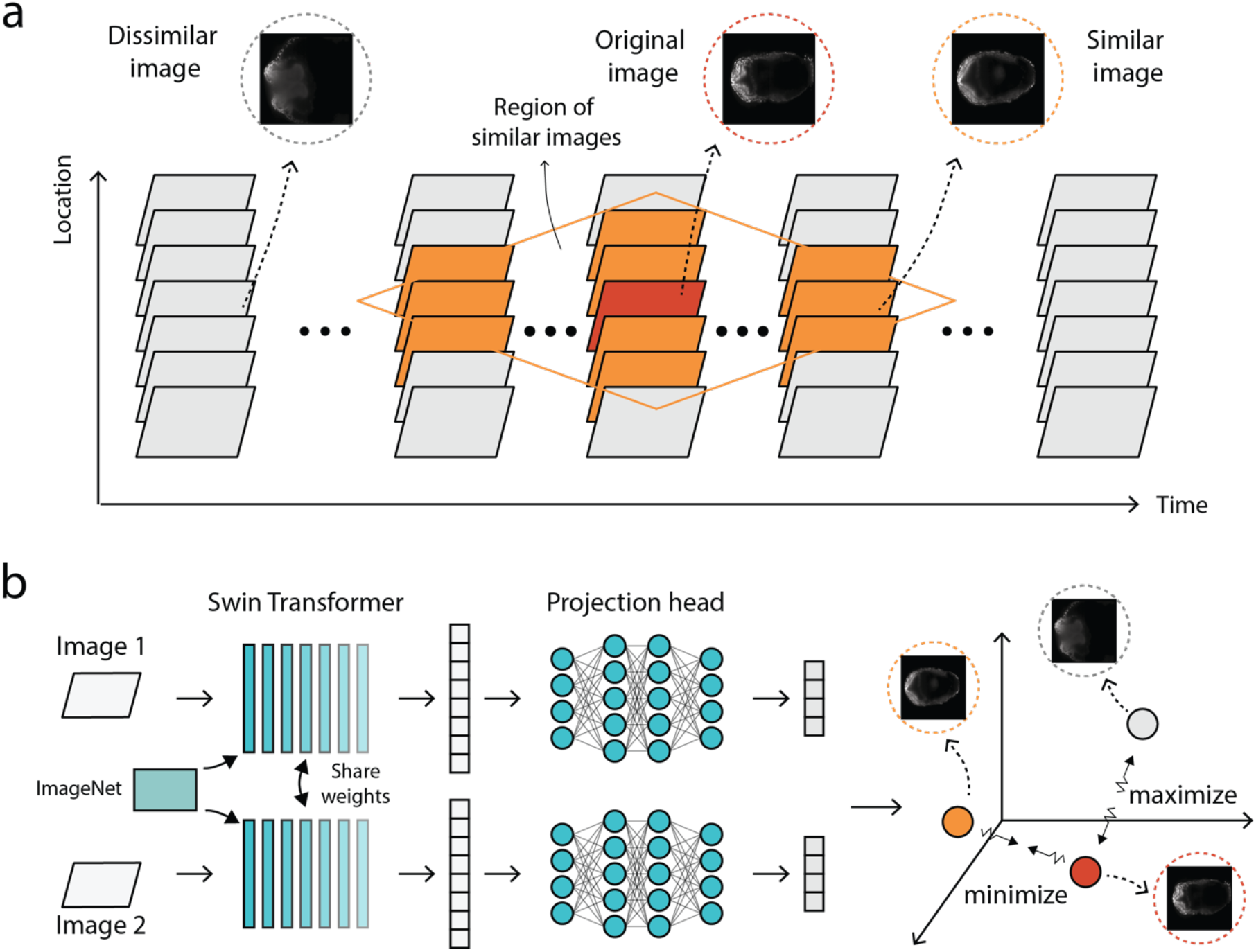
Dataset construction and contrastive training for STERN. **a**, Schematic of dataset construction from embryo imaging data (Embryos A, B and C are used for training). For an original image slice (red), candidate slices are grouped by spatial and/or temporal proximity (location and time). Slices within a local neighbourhood (orange) are treated as similar candidates (positive pairs), whereas distant slices are treated as dissimilar candidates (negative pairs). Representative similar and dissimilar examples are shown. **b**, Model construction and contrastive training. Pairs of image slices are processed by a Siamese Swin Transformer backbone (initialized from ImageNet pre-training) with shared weights, followed by a projection head. A contrastive objective is used to learn an embedding space in which similar pairs are pulled together (minimized distance) and dissimilar pairs are pushed apart (maximized distance).

**Extended Data Fig. 2.**
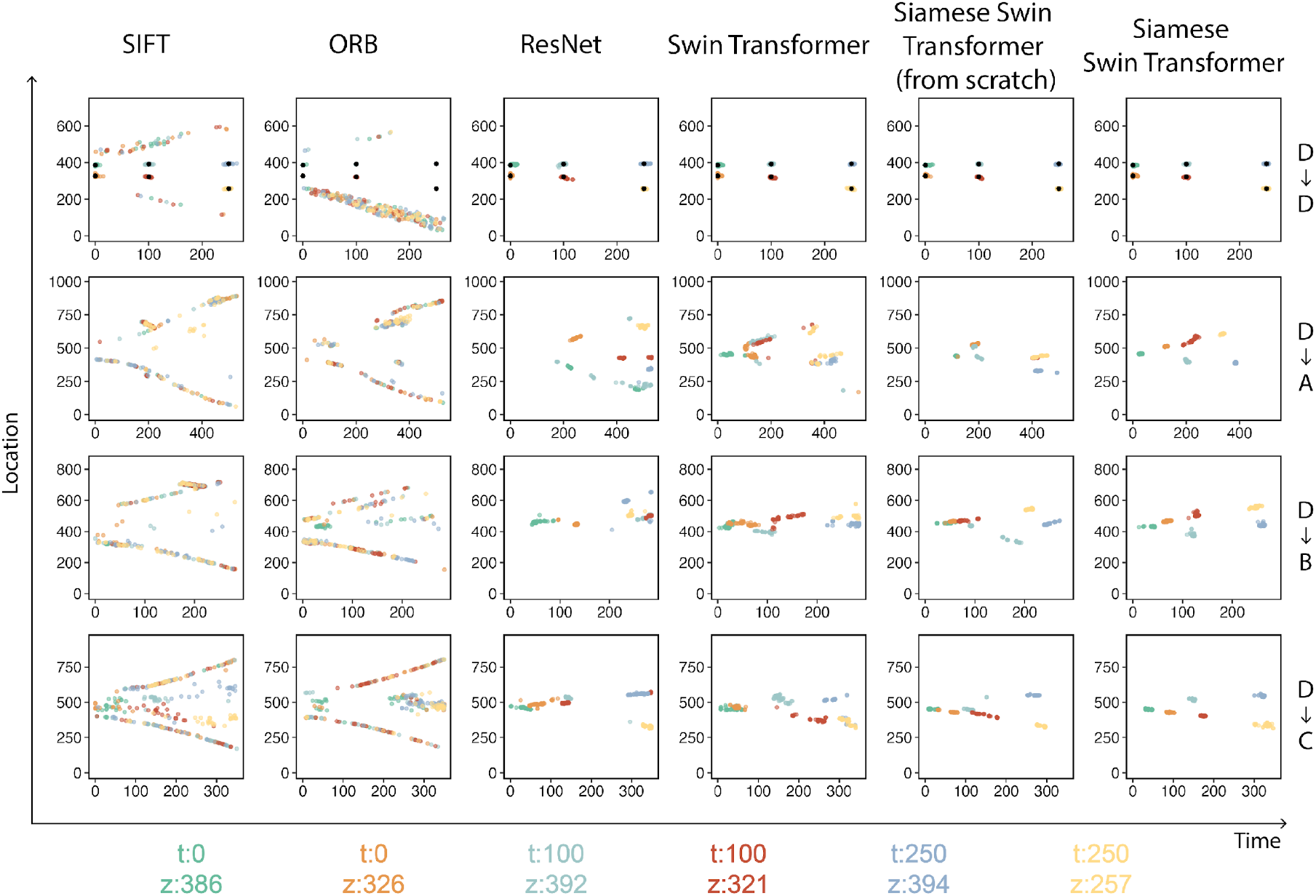
Comparative analysis of cross-embryo matching using six query slices with varying time points and locations. Two traditional methods, SIFT and ORB, are compared against four machine learning methods: two pretrained models (ResNet and Swin Transformer), a Siamese Swin Transformer trained from scratch, and a Siamese Swin Transformer fine-tuned from the pretrained Swin Transformer (shown column-wise). “D->D” denotes matching query slices against Embryo D itself, while “D->A/B/C” represents matching query slices against Embryos A, B, and C, respectively. Each line in a sub-plot shows the spatial and temporal distributions of the top 50 matching slices for each query slice. In the “D->D” case, the location of the query slice is marked by a black dot as a reference anchor point.

**Extended Data Fig. 3.**
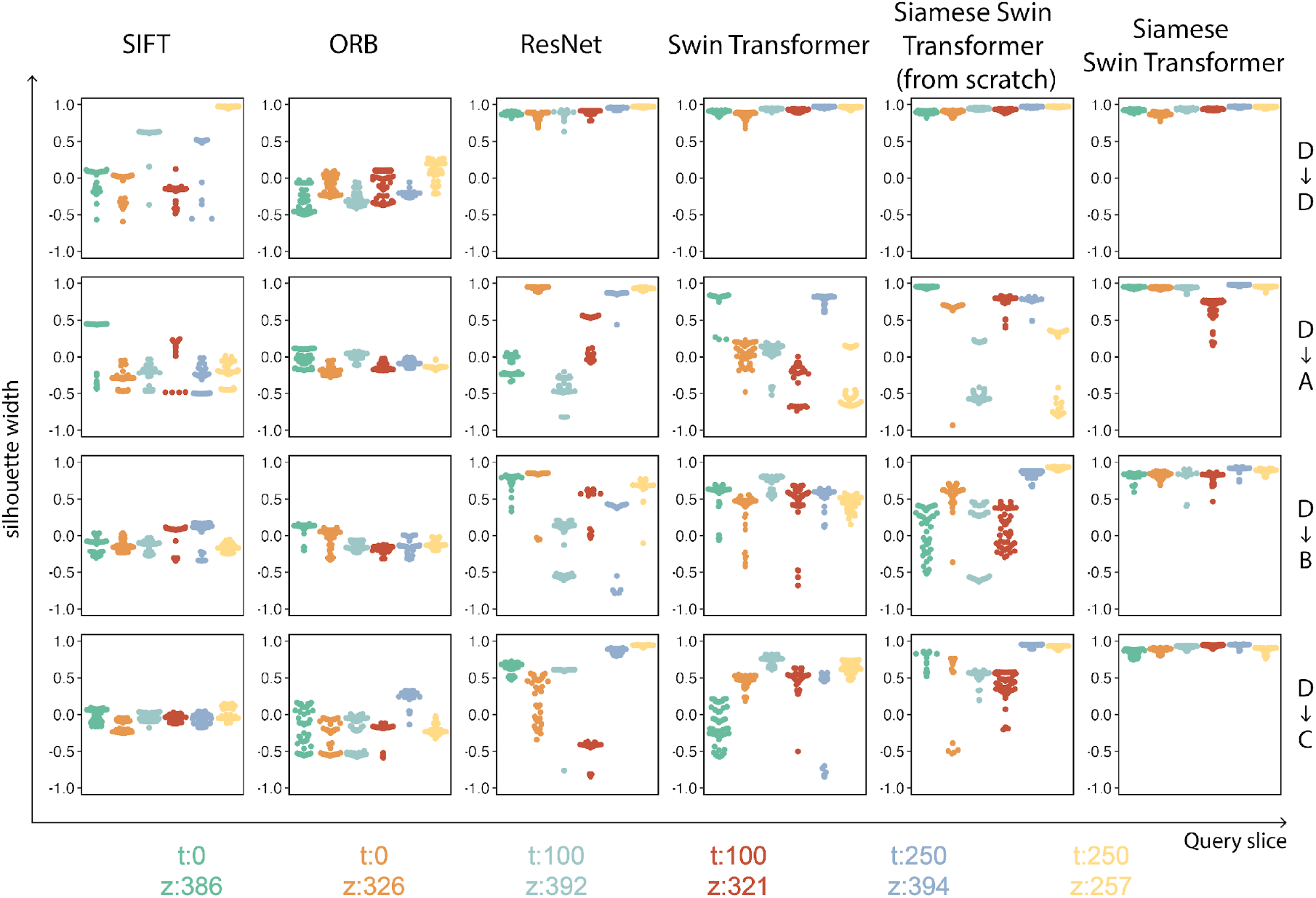
Quantitative analysis of cross-embryo matching using Silhouette width. Each box plot represents the Silhouette width values calculated for the top 50 matched points from the scatter plots in Extended Data Fig. 2, categorized by the six query slices.

**Extended Data Fig. 4.**
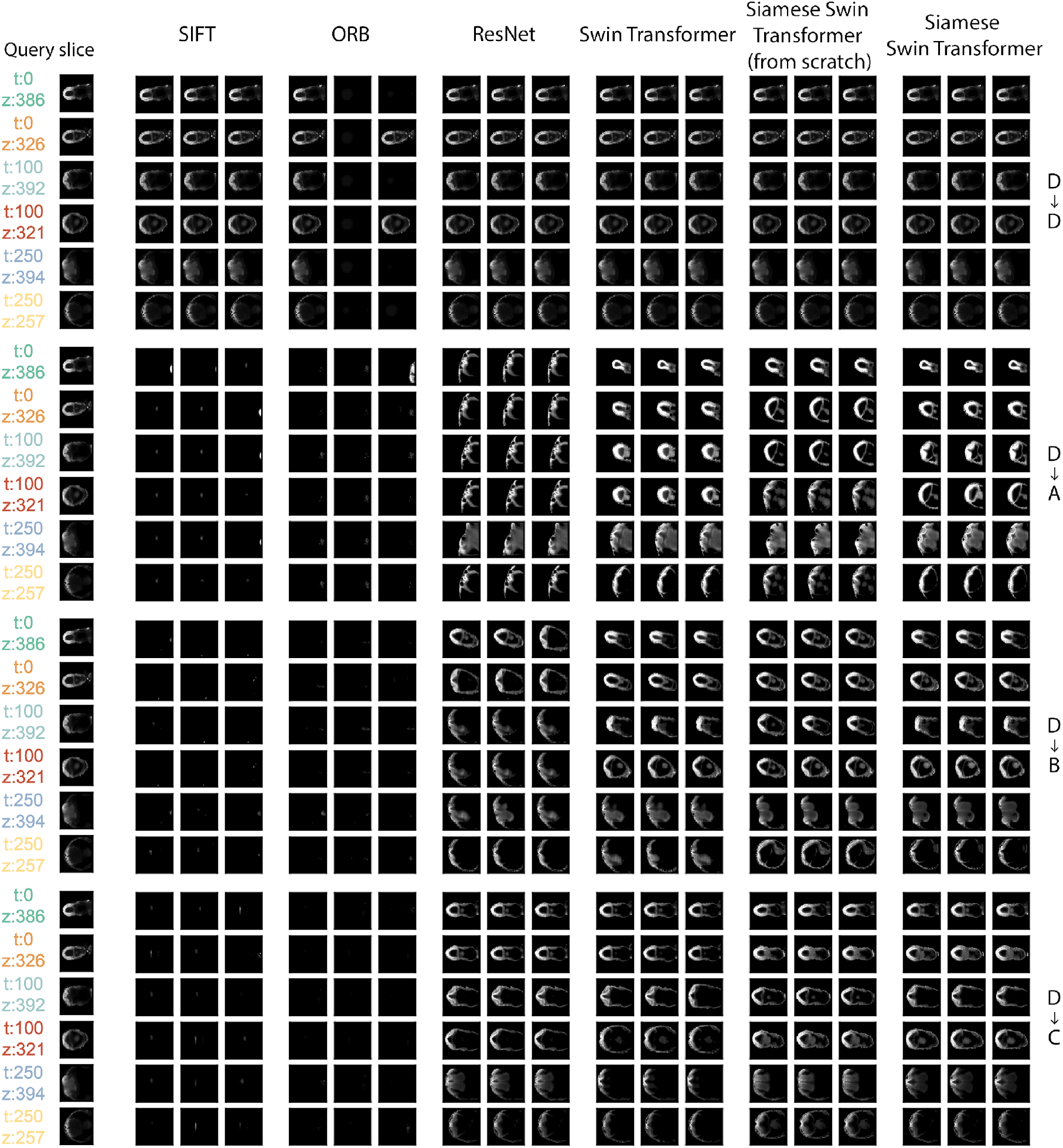
Comparative analysis of the top three matching results from six methods for cross-embryo matching. Both self-embryo matching (D->D) and cross-embryo matching (D->A/B/C) results are presented for each of the six query slices. The leftmost column lists the six query slices, while the subsequent columns showcase the top three matches obtained by each model.

**Extended Data Fig. 5.**
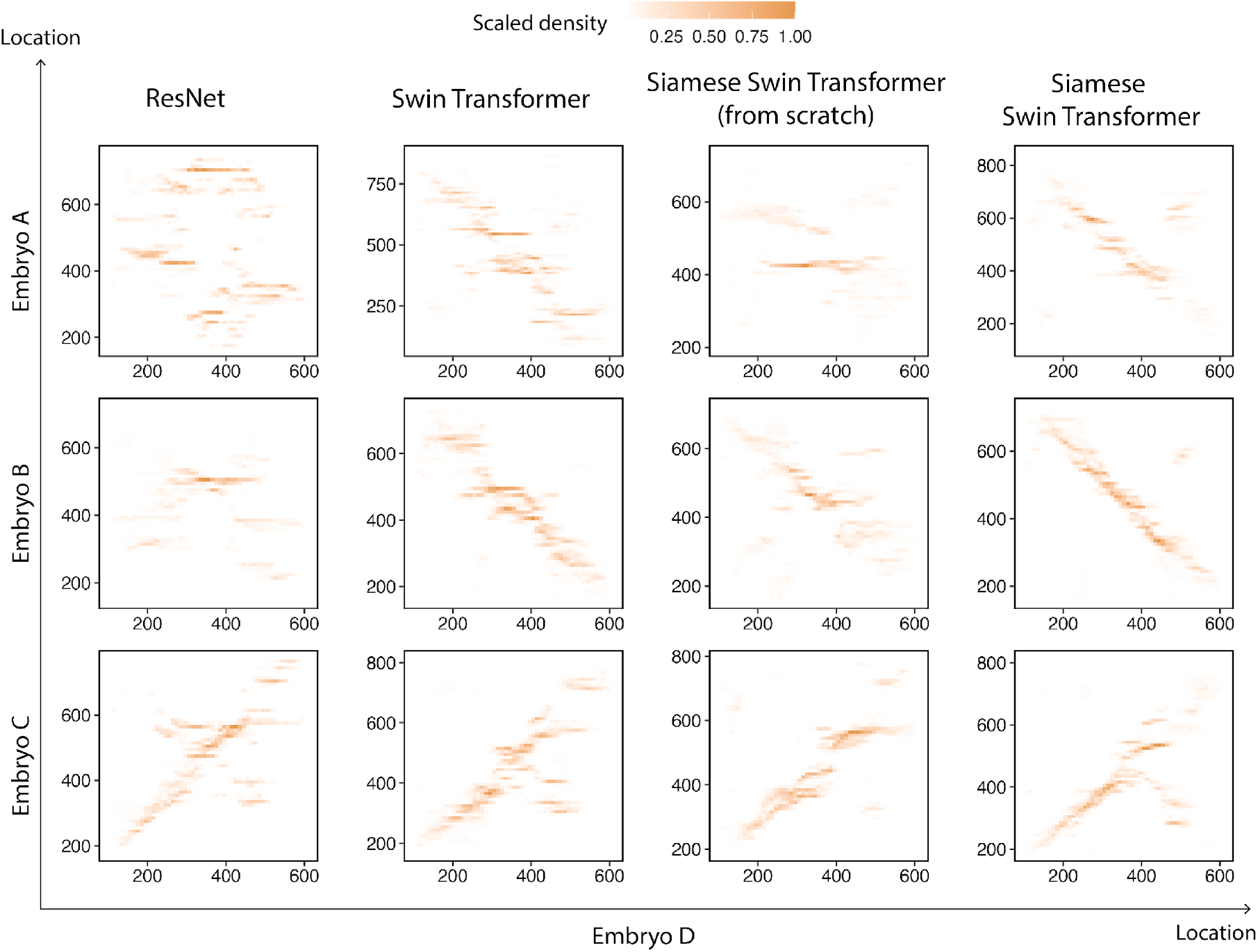
Quantitative analysis of spatial relationships between embryos. The sub-plots visualize the same slice pairs, generated in Extended Data Fig. 6, to explore spatial relationships between Embryo D and Embryos A, B, and C. The x-axis represents the location of query slices in Embryo D, while the y-axis displays the location of the top-matched slices in the other embryos (shown row-wise). Colour intensity follows the same scale as in Extended Data Fig. 6, where it indicates the density of points in 10 by 10 bins, scaled to a maximum of 1.

**Extended Data Fig. 6.**
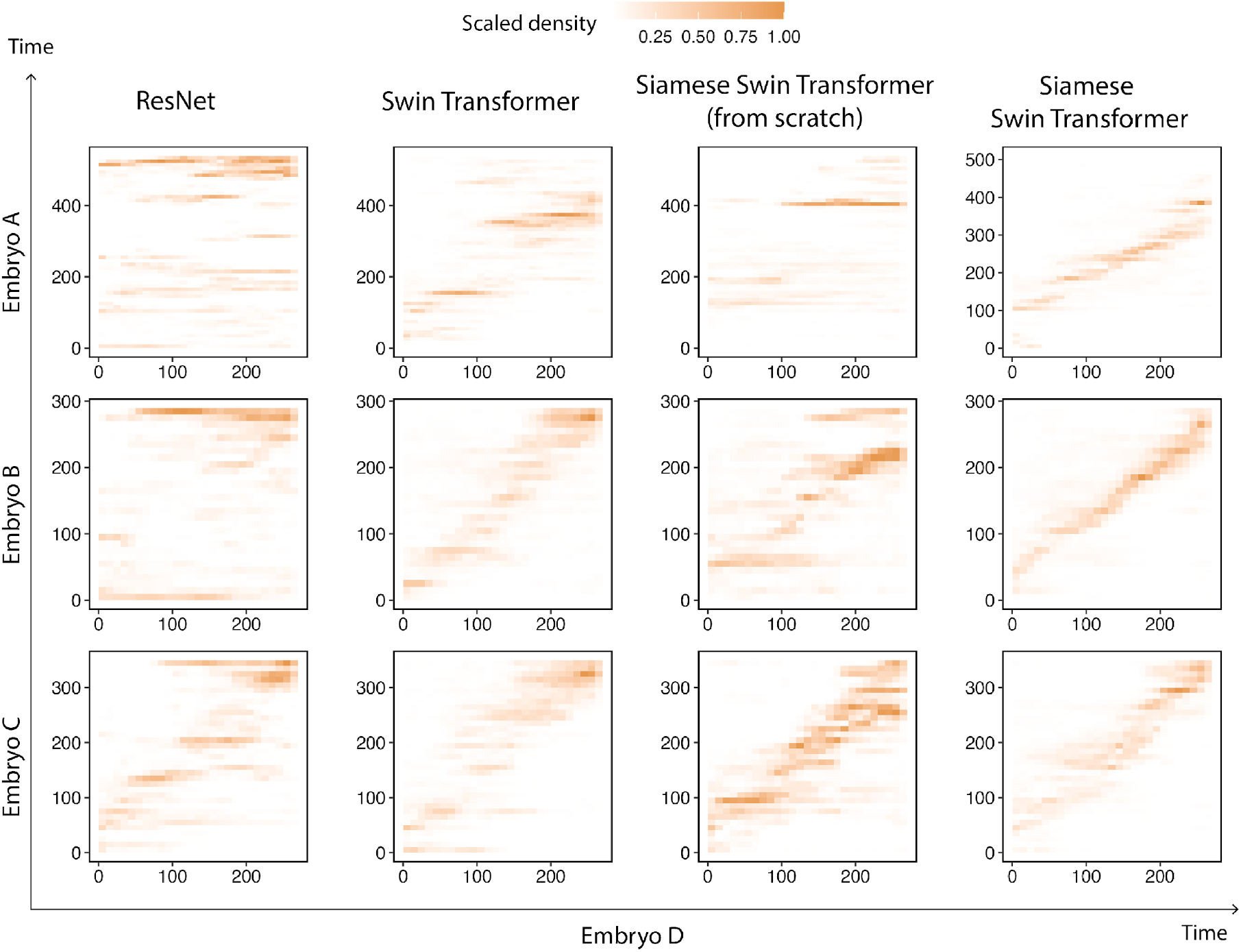
Quantitative analysis of temporal relationships between embryos. For all non-empty image slices (referred to as query slices) from Embryo D, the closest matching slices in Embryos A, B, and C were identified using four deep learning-based methods (shown column-wise), resulting in pairs of query slices and their top-matched slices. The sub-plots visualize these slice pairs to explore temporal relationships between Embryo D and the other embryos, where the x-axis represents the time points of query slices in Embryo D and the y-axis displays the time points of the top-matched slices in Embryos A, B, and C (shown row-wise). Colour intensity indicates the density of points within a corresponding 10 by 10 bin, scaled to a maximum of 1.

**Extended Data Fig. 7.**
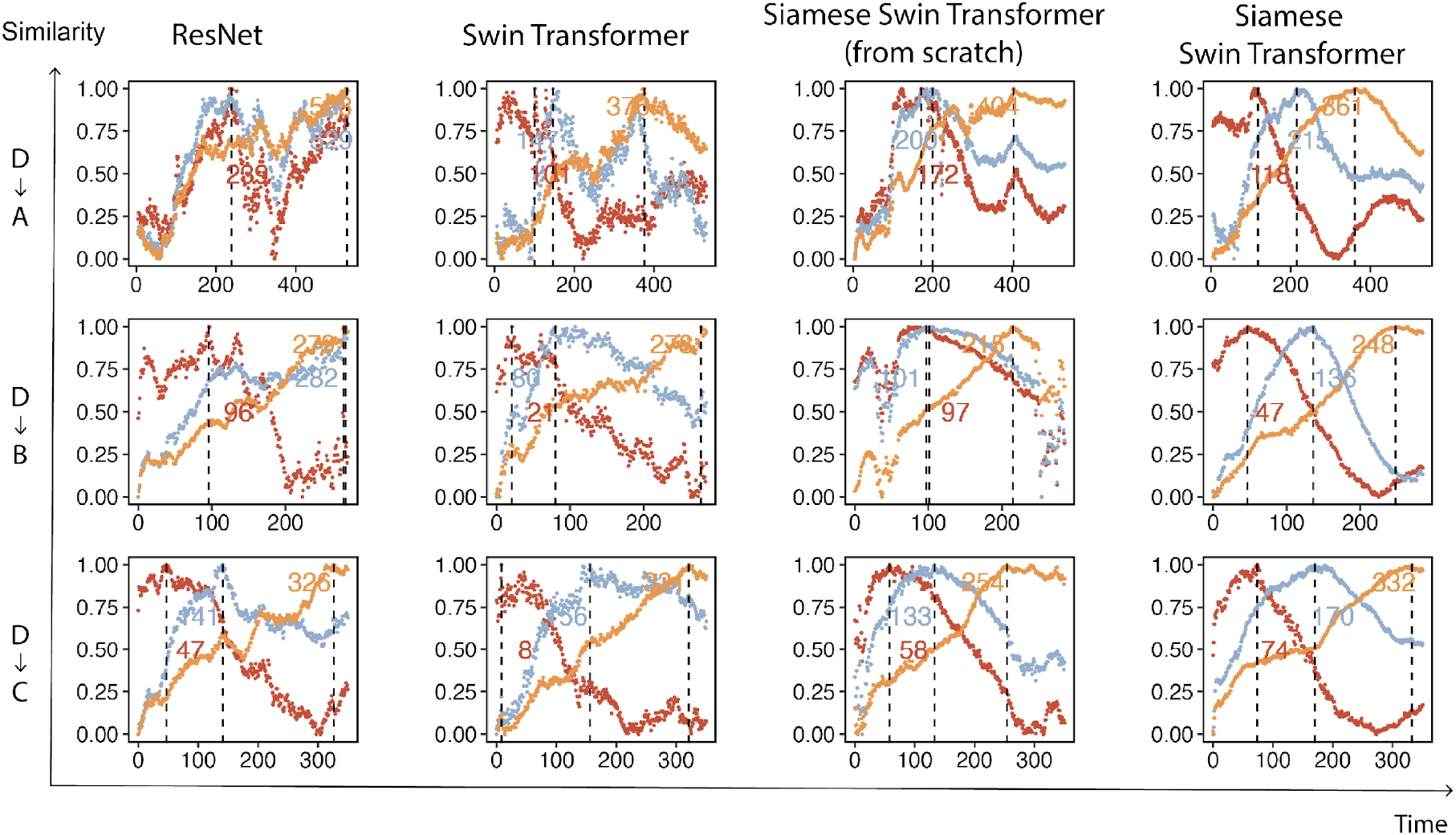
Comparative analysis of deep learning methods on 3D cross-embryo matching. The results of each model are shown column-wise. The distribution of similarities between the three query embryos in Embryo D and Embryos A, B, and C, respectively (“D->A/B/C”), is denoted by three coloured lines. For each query embryo, a vertical dashed line indicates the best-matched embryo, with the corresponding time points marked adjacent to the dashed line.

**Extended Data Fig. 8.**
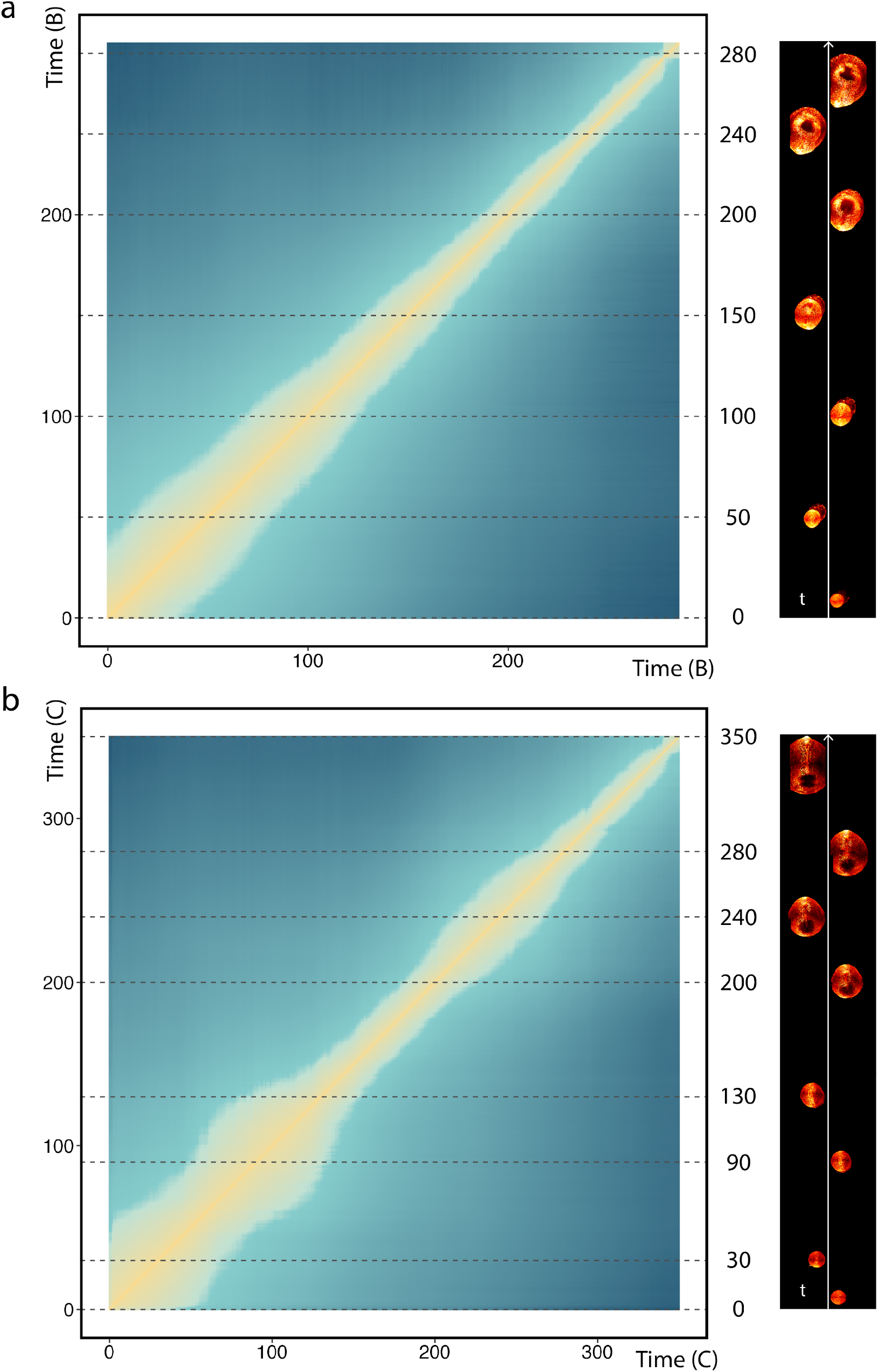
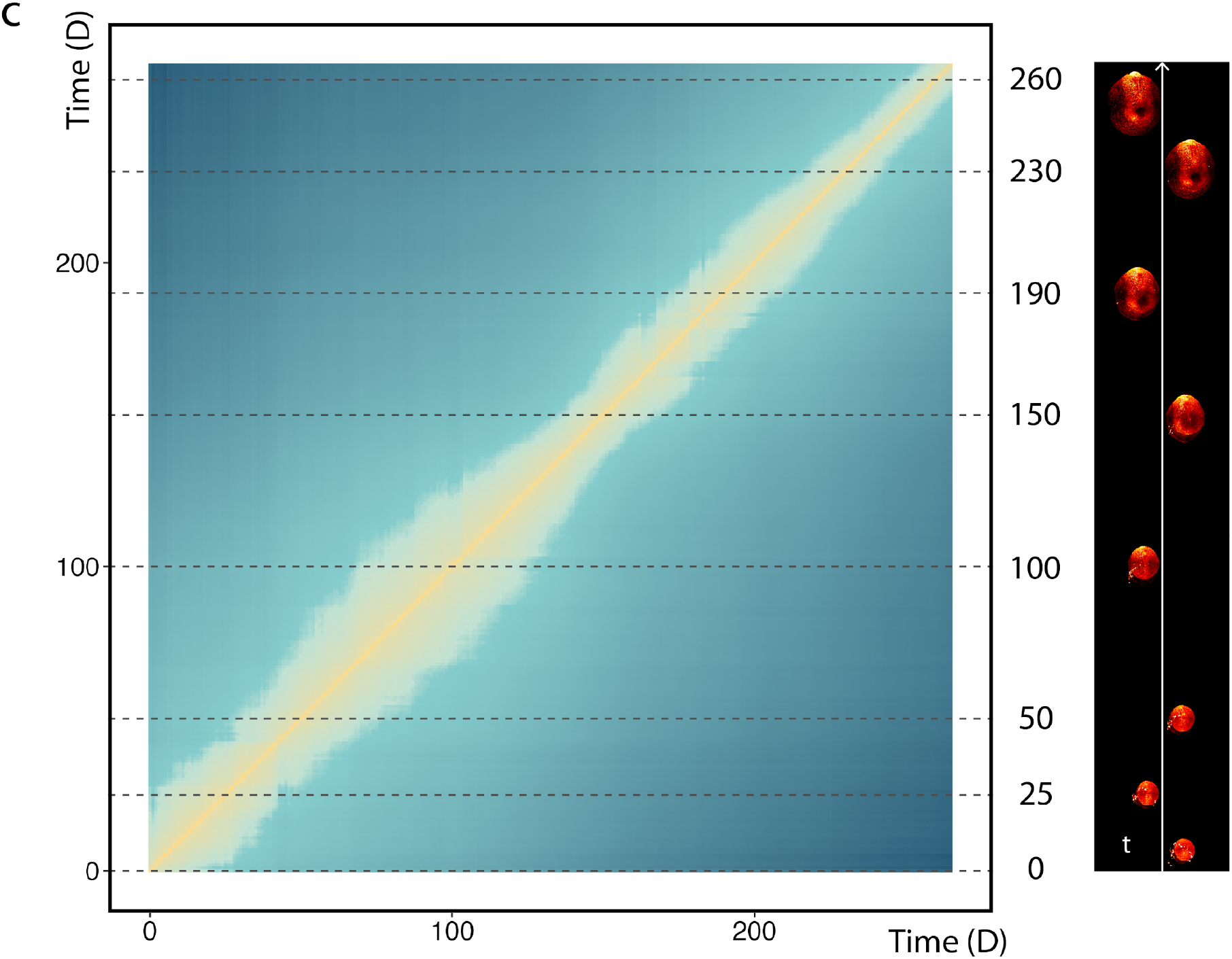
Similarity matrices for embryo pairs from each of embryos B, C and D. Similarities between embryo pairs at all time points within (**a**) B, (**b**) C, and (**c**) D were calculated in the same manner as in Fig. 3a. Horizontal dashed lines highlight regions of interest in each matrix, with the corresponding embryos displayed on the right.

**Extended Data Fig. 9.**
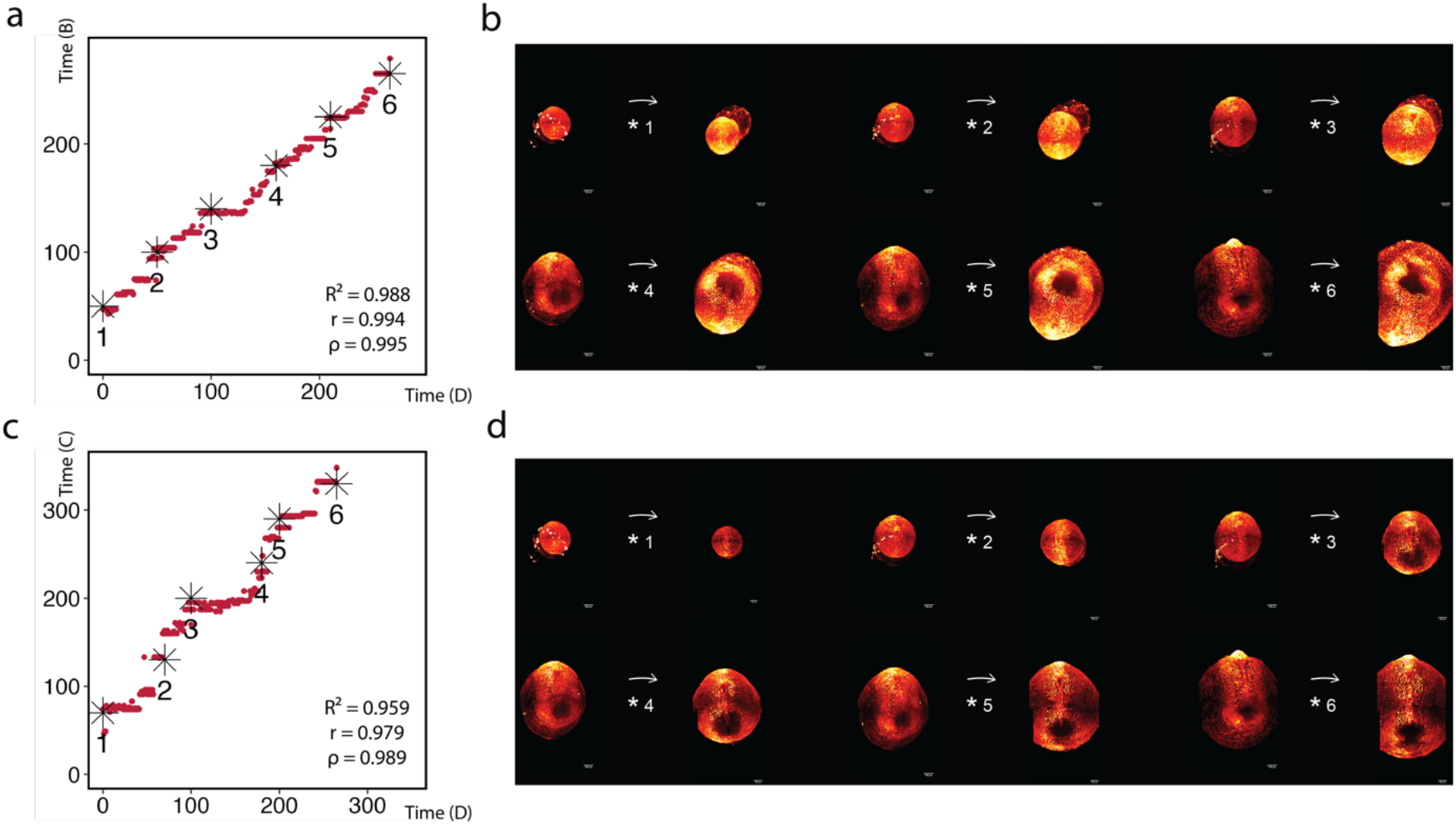
Cross-embryo 3D temporal alignment between Embryo D and Embryos B and C. **a**, Temporal alignment between Embryos D and B. The scatter plot shows the best-matched developmental time point in Embryo B for each query time point in Embryo D. Representative matched pairs are marked as *1–*6. Pearson correlation, Spearman rank correlation and the coefficient of determination from a linear fit are shown. **b**, Representative matched 3D embryo volumes corresponding to the pairs marked in a. For each pair, the query embryo (Embryo D) is shown on the left and the matched reference embryo (Embryo B) on the right. **c**, Temporal alignment between Embryos D and C. The scatter plot shows the best-matched developmental time point in Embryo C for each query time point in Embryo D. Representative matched pairs are marked as *1–*6. Pearson correlation, Spearman rank correlation and the coefficient of determination from a linear fit are shown. **d**, Representative matched 3D embryo volumes corresponding to the pairs marked in c. For each pair, the query embryo (Embryo D) is shown on the left and the matched reference embryo (Embryo C) on the right.

**Extended Data Fig. 10.**
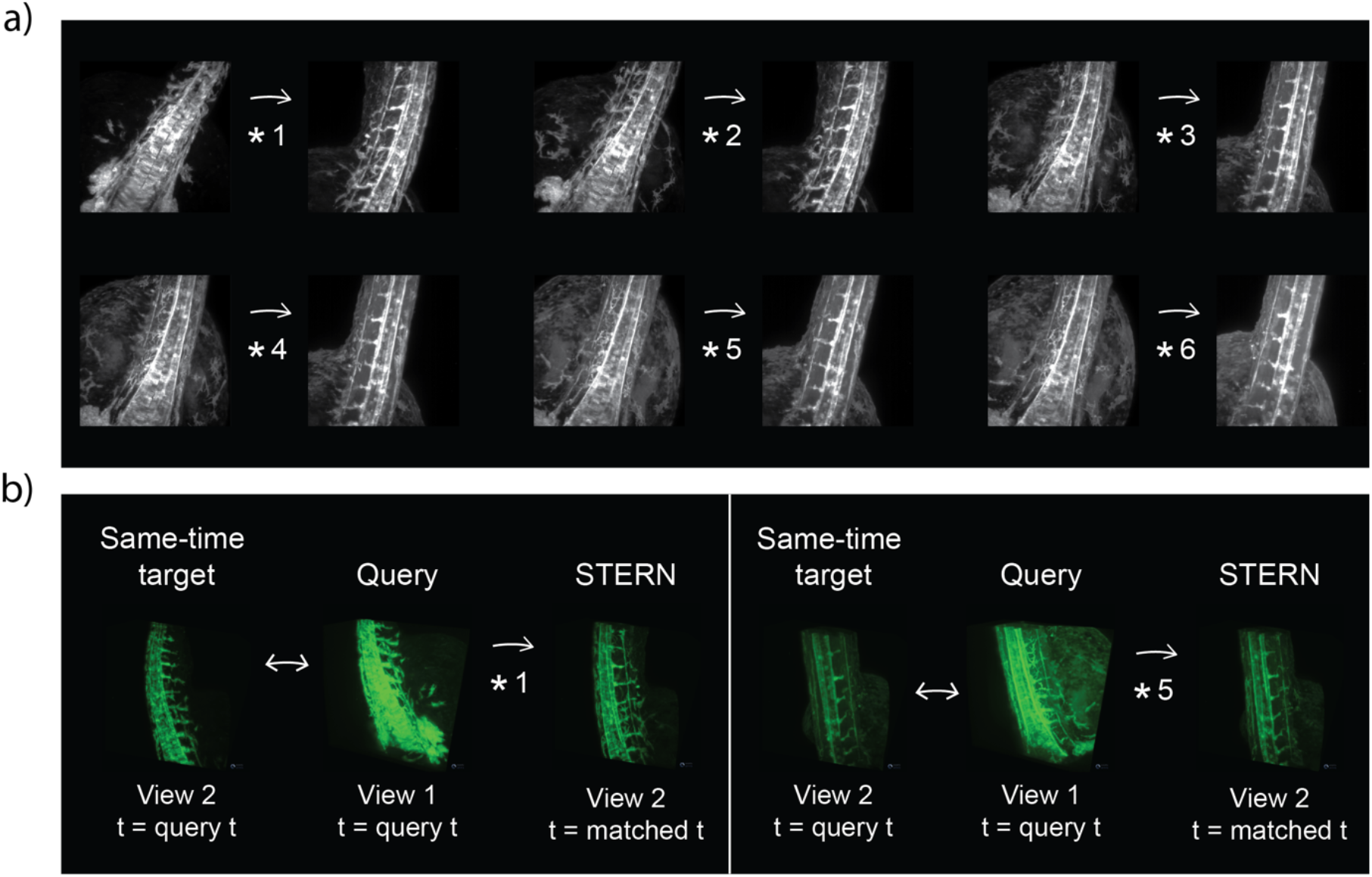
Visual validation of zebrafish cross-view matched states. **a**, Maximum-intensity projection views of the six representative zebrafish cross-view matched states marked in Fig. 3h and shown as volume-rendered views in Fig. 3i. Projections were generated from the corresponding z-stacks along the acquisition direction, without manual adjustment of the viewing angle. For each pair, the query state from view 1 is shown on the left and the STERN-selected matched state from view 2 is shown on the right. **b**, Comparison of same-time and STERN-selected target states for two representative off-diagonal matches (*1 and *5). Viewing angles were kept fixed across time points after choosing an initial comparable perspective. For each example, the same-time target state from view 2 is shown at the query time point, the query state from view 1 is shown at the same time point, and the STERN-selected target state from view 2 is shown at the matched time point.

**Extended Data Fig. 11.**
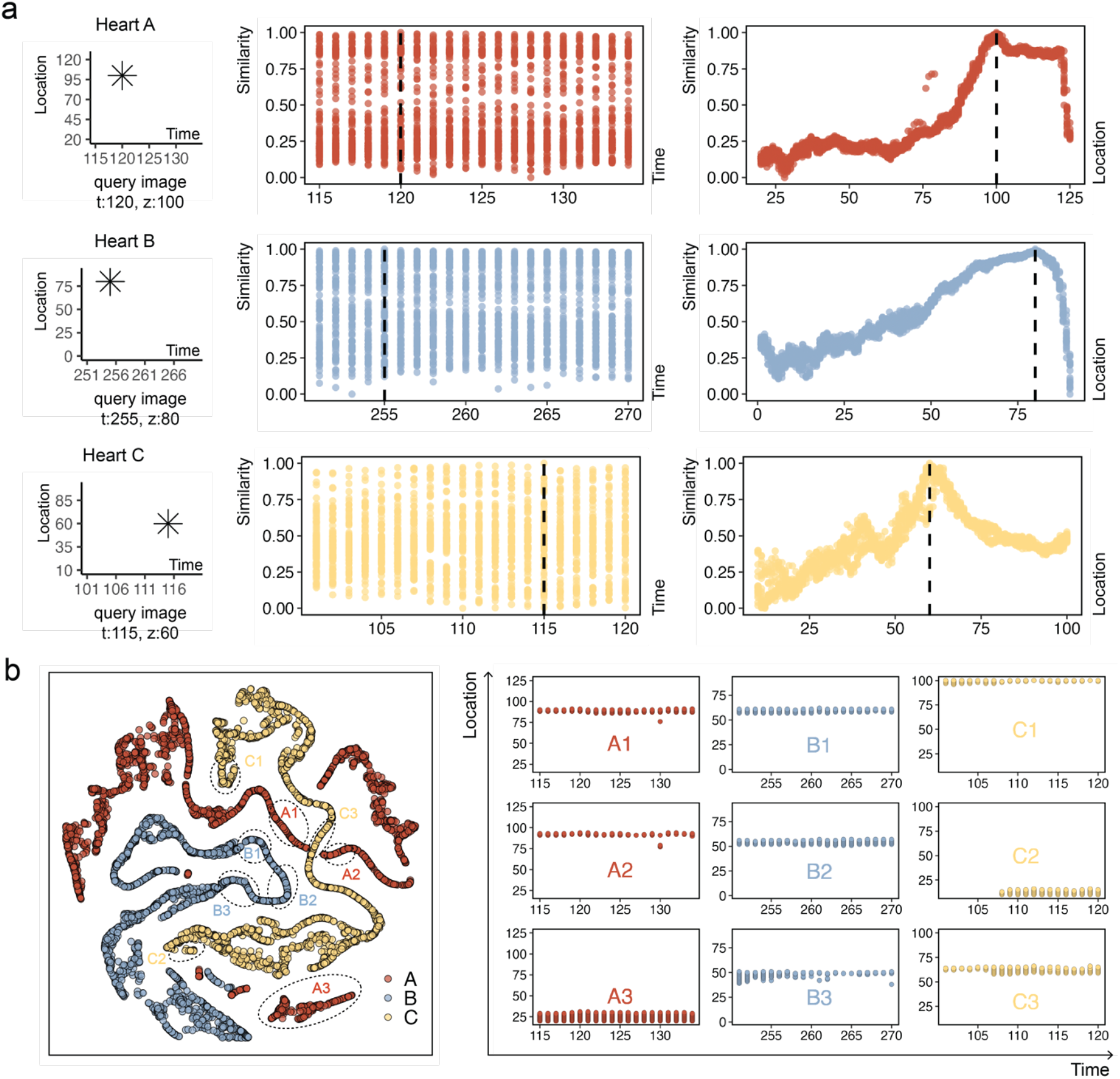
Analysis of real-time developing mouse hearts at the 2D image level. **a**, Similarity plots for three selected query images (t:120z:100 from Heart A, t:255z:80 from Heart B and t:115z:60 from Heart C) are categorized into three rows, distinguished by corresponding colours. Each row includes similarity trends across time (left) and location (right). The black vertical dotted line in each plot indicates the original position of the query image in either time or location. **b**, Three heart datasets, denoted as Hearts A, B, and C (highlighted by different colours), represent distinct developmental stages of 3D real-time developing mouse hearts. The datasets were sliced along the original observation direction (z-axis, similar to that of above embryos). Empty slices were removed before downstream analysis. Our model extracted features from slices in all three datasets, and dimensionality reduction was applied to visualize them in a 2D space (left panel). For each dataset, three interesting distributions (A1-A3, B1-B3, and C1-C3) were selected, and slices from these areas were projected to their original spatial and temporal spaces (right panel).

**Extended Data Fig. 12.**
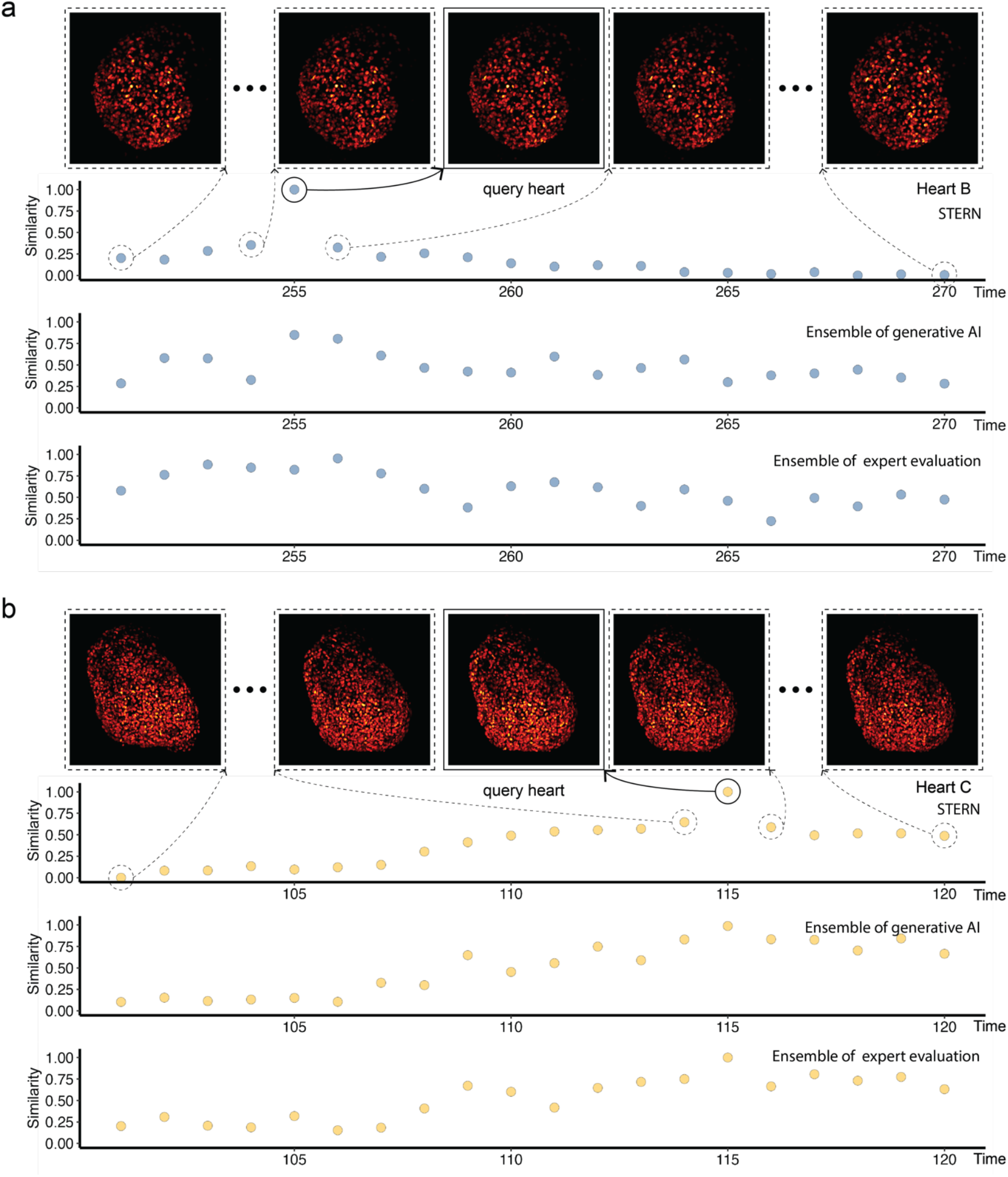
Temporal similarity trajectories across additional heart datasets. **a**, Comparison of temporal similarity trajectories for Heart B using a query volume at time point t = 255. **b**, Comparison of temporal similarity trajectories for Heart C using a query volume at time point t = 115. For each dataset, the three lower panels show normalized similarity trajectories produced by STERN (top), the multimodal AI ensemble (middle), and the expert ensemble (bottom), respectively, when matching the query heart volume against all time points from the same dataset. Higher scores indicate greater similarity to the query. The image panels show representative reference volumes sampled from distinct regions of the corresponding STERN trajectories, including highly similar and weakly similar matches relative to the query heart volume. Dashed lines connect selected trajectory positions to their corresponding representative images. All scores were normalized to fall between 0 and 1.

## Data availability

Mouse embryo imaging data are publicly available through the Image Data Resource (IDR) repository (https://idr.openmicroscopy.org/webclient/?show=project-502). Mouse heart imaging data are publicly available through Figshare (https://figshare.com/projects/Long-term_live_imaging_of_mouse_embryonic_heart/74532). Zebrafish imaging data generated in this study will be deposited in a public repository upon publication and are available upon reasonable request to the corresponding authors.

## Code availability

The source code underlying STERN will be made publicly available upon publication. Prior to public release, the code is available upon reasonable request to the corresponding authors.

## Acknowledgments

The authors thank James Sharpe (EMBL Barcelona) for valuable discussions and scientific insights. J.W. was a recipient of the Marie Skłodowska-Curie Postdoctoral Fellowship (Grant Agreement No. 101067151), the EMBL Interdisciplinary Postdoctoral (EIPOD) Fellowship and the EMBO Non-Stipendiary Fellowship (EMBO ALTF 400-2022), and was a Junior Research Fellow at Wolfson College, University of Cambridge, UK. J.W. acknowledges support from the Faculty of Science start-up funding at the University of Bath. B.D. acknowledges support from the National Institute of General Medical Sciences of the National Institutes of Health under Award Number R35GM157082, from the CIFAR Azrieli Global Scholars program, and from the CIFAR MacMillan Multiscale Human.

