## Supplementary Information for "A quantitative coordinate system for developmental dynamics"

Supplementary File

### Supplementary Methods

#### Selection of positive samples

Here, each positive sample denotes a labelled image-pair used for contrastive training, in which the two image slices are expected to have similar morphology. Positive samples were generated through five steps.

***1. Defining embryo bounds and sections.*** We began by traversing all time points containing non-empty z-slices for each embryo. For a given time point $t_i$, we established the lower bound $z_L^{t_i}$ and upper bound $z_U^{t_i}$ before dividing the z-slices into four sections. Starting from section 1 at time $t_i$, we examined the z-slices individually. Let $z_{1,k}^{t_i}$ represent the z-coordinate of the $k_{th}$ z-slice in section 1 at time $t_i$, arranged in ascending order such that $z_{1,k}^{t_i} < z_{1,k+1}^{t_i}$. We let the z-slice at $z_{1,k}^{t_i}$ be an original slice.

***2. Selecting candidate positive slices within the same time point.*** Let $z_{lub}^{t_i}$ and $z_{glb}^{t_i}$ denote the least upper bound and greatest lower bound that capture z-slices similar to $z_{1,k}^{t_i}$ at time point $t_i$. The potential z-slices for constructing positive samples with respect to $z_{1,k}^{t_i}$ are between $z_{glb}^{t_i}$ and $z_{lub}^{t_i}$. $z_{glb}^{t_i}$ is calculated as $z_{glb}^{t_i} = z_{1,k}^{t_i} - \left(z_{lub}^{t_i} - z_{1,k}^{t_i}\right)$ so that it is symmetric to $z_{lub}^{t_i}$ with respect to $z_{1,k}^{t_i}$, however we excluded z-slices between $z_{glb}^{t_i}$ and $z_{1,k}^{t_i}$, to avoid duplicating positive samples during traversal. The remaining z-slices between $z_{1,k+1}^{t_i}$ and $z_{lub}^{t_i}$ contribute to the final positive sample set for the original slice at $z_{1,k}^{t_i}$.

The value of $z_{lub}^{t_i}$ is determined by adjusting $z_{3,1}^{t_i}$, the z-coordinate of the first z-slice more than one section away from $z_{1,k}^{t_i}$, using a manually determined factor called $z_{shrink}$. The values of this factor, which varies for each embryo, are detailed in Supplementary Table S2. The value of $z_{lub}^{t_i}$ is calculated as $z_{lub}^{t_i} = z_{1,k}^{t_i} + (z_{3,1}^{t_i} - z_{1,k}^{t_i})/z_{shrink}$.

***3. Excluding ambiguous spatial slices.*** Z-slices between $z_{lub}^{t_i}$ and $z_{3,1}^{t_i}$ were assigned to the ambiguous class and excluded from positive sample selection to reduce the chance of assigning visually uncertain samples to the positive sample set.

***4. Extending positive sample selection across nearby time points.*** We next extended positive sample selection to subsequent time points, as slices at a fixed z-coordinate can remain morphologically similar over a short developmental interval. Time points before $t_i$ are neglected to avoid duplicating positive samples. Let $t_A$ represent the number of time points it will take for a z-slice at a fixed z-coordinate to manifest apparent changes. STERN training used a time-dependent sampling parameterization in which $t_A$ was specified separately for two developmental windows for each embryo ($t_{A,1}$ and $t_{A,2}$; Supplementary Table S2). The start points of the two windows were defined by the embryo-specific values $t_{start,1}$ and $t_{start,2}$ listed in Supplementary Table S2. For a given original slice at time point $t_i$, the value of $t_A$ was selected according to the developmental window containing $t_i$. For notational clarity, the equations below use $t_A$ as a generic symbol; in implementation, $t_A$ denotes the appropriate first- or second-window value assigned according to $t_i$.

Let $t_P$ represent the number of time points that a z-slice at a fixed z-coordinate stays in a similar appearance. The value of $t_P$ is determined by adjusting the selected value of $t_A$ using the manually determined temporal shrink factor $t_{shrink}$ (Supplementary Table S2). The value of $t_P$ is calculated as $t_P = t_A / t_{shrink}$. As such, we let the time points between $t_i$ and $t_i + t_P$ be the time points where further z-slices will be considered for positive samples, and the time points between $t_i + t_P$ and $t_i + t_A$ are no longer considered because they are considered as ambiguous.

Having determined the available time points for generating further positive samples, we then needed to determine the appropriate upper bounds for those time points to determine the z-slices for positive samples. Let $t_j$ be a time point between $t_i$ and $t_i + t_P$. We infer the $z_{lub}^{t_j}$ of z-slices similar to the $z_{1,k}^{t_i}$ at $t_j$ using linear interpolation between two points: $(t_i, z_{lub}^{t_i})$ and $(t_i + t_P, z_{1,k}^{t_i})$. The corresponding greatest lower bound $z_{glb}^{t_j}$ is thus calculated similarly as aforementioned, $z_{glb}^{t_j} = z_{1,k}^{t_i} - \left(z_{lub}^{t_j} - z_{1,k}^{t_i}\right)$. In cases where $z_{glb}$ is less than $z_L$, we set $z_{glb}$ to be equal to $z_L^{t_j}$. Therefore, for the original slice at $z_{1,k}^{t_i}$, z-slices between $z_{glb}^{t_j}$ to $z_{lub}^{t_j}$ were assigned to the positive sample set at time point $t_j$ By repeating the same operation for all time points between $t_i$ and $t_i + t_P$, we selected z-slices that are used for constructing positive samples for the original slice at $z_{1,k}^{t_i}$.

***5. Applying the procedure to other sections and final filtering.*** The selection of z-slices for positive samples for original slices in section 2 followed a similar procedure, except that $z_{3,1}^{t_i}$ was replaced by $z_{4,1}^{t_i}$ when computing $z_{lub}^{t_i}$, with the remaining values adjusted accordingly. For an original slice in section 3, $z_{3,k}^{t_i}$ at time point $t_i$, the situation is slightly different. We computed the $z_{glb}^{t_i}$ first and then used $z_{glb}^{t_i}$ to determine the value of $z_{lub}^{t_i}$ using its symmetric property. Since the adjacent section is section 2, we considered the z-slices above the minimum z-coordinate of section 2 as candidate positive samples. Accounting for the ambiguous class, $z_{glb}^{t_i}$ is computed as $z_{glb}^{t_i} = z_{3,k}^{t_i} - (z_{3,k}^{t_i} - z_{2,1}^{t_i})/z_{shrink}$ and $z_{lub}^{t_i}$ is computed as $z_{lub}^{t_i} = z_{3,k}^{t_i} + (z_{3,k}^{t_i} - z_{glb}^{t_i})$. If $z_{lub}^{t_i}$ exceeds $z_U^{t_i}$, we set $z_{lub}^{t_i}$ to be equal to $z_U^{t_i}$. Once we have determined those two values, the selection of positive samples followed the same approach as before.

The process for generating positive samples of an original slice $z_{4,k}^{t_i}$ in section 4 at time point $t_i$ is similar to the original slices in section 3, except that we calculated $z_{glb}^{t_i}$ as follows: $z_{glb}^{t_i} = z_{4,k}^{t_i} - (z_{4,k}^{t_i} - z_{3,1}^{t_i})/z_{shrink}$ by considering z-slices above the minimum z-coordinate of section 3 as candidate positive samples.

The above process describes the generation of positive samples for original slices of 4 sections at $t_i$. In the actual sample selection, we observed a substantial number of incorrect positive samples, in which the two selected images did not appear morphologically similar. Upon investigation, we realized that the second slices generated from original slices in section 1 and section 4 were major contributors to this issue. Section 1 and section 4 contain many edge slices that exhibit substantial variations in comparison to the image slices at the central location of the embryo (section 2 and 3). Consequently, original slices from sections 1 and 4 were excluded from the final positive sample construction.

**Selection of negative samples**

Here, each negative sample denotes a labelled image-pair used for contrastive training, in which the two image slices are expected to be morphologically dissimilar. Negative samples were selected through four steps.

***1. Defining spatially separated candidate negative samples.*** Like the selection of positive samples, we began by traversing all time points containing non-empty z-slices for each embryo. For a given time point $t_i$, we established the lower bound $z_L^{t_i}$ and upper bound $z_U^{t_i}$ to divide the z-slices into four sections. Starting from section 1 at time $t_i$, we examined the z-slices individually. Let $z_{1,k}^{t_i}$ represent the z-coordinate of the $k_{th}$ z-slice in section 1 at time $t_i$, arranged in ascending order such that $z_{1,k}^{t_i} < z_{1,k+1}^{t_i}$ . We let the z-slice at $z_{1,k}^{t_i}$ be an original slice and considered z-slices that are one section away from it as candidate negative samples. Let $z_{glb}^{t_i}$ be the greatest lower bound of z-slices that are considered as not similar to the original slice at $z_{1,k}^{t_i}$, the $z_{glb}^{t_i}$ is set to be $z_{3,1}^{t_i}$ and this serves a different purpose compared to selecting positive samples. As such, z-slices between $z_{glb}^{t_i}$ and $z_U^{t_i}$ are assigned to the candidate negative sample set for the original slice $z_{1,k}^{t_i}$ at $t_i$. Time points before $t_i$ are not considered to avoid repeating negative samples during traversal. As such, time points after $t_i$ are considered for negative sample selection and we separated them into two cases:

***2. Extending negative sample selection across time.*** Case 1: Let $t_N$ represents the number of time points required for a z-slice at a fixed z-coordinate to become visibly dissimilar. As with $t_A$, $t_N$ was specified separately for the two developmental windows in the time-dependent sampling parameterization ($t_{N,1}$ $and$ $t_{N,2}$; Supplementary Table S2) , with the appropriate value selected according to the original time point $t_i$. If a time point $t_j$ is beyond $t_i + t_N$, then z-slices between $z_L^{t_j}$ and $z_U^{t_j}$ at time $t_j$ are considered as candidate negative samples with respect to original slice $z_{1,k}^{t_i}$.

Case 2: If a time point $t_j$ is between $t_i$ and $t_i + t_N$, we estimate the $z_{glb}^{t_j}$ of the z-slices dissimilar to $z_{1,k}^{t_i}$ using linear interpolation based on two points: $(t_i, z_{3,1}^{t_i})$ and $(t_i + t_N, z_{1,k}^{t_i})$. Consequently, z-slices between $z_{glb}^{t_j}$ and $z_U^{t_j}$ are considered as candidate negative samples with respect to original slice $z_{1,k}^{t_i}$.

***3. Applying the procedure to other sections.*** Combining cases 1 and 2 allows us to determine the candidate negative samples for original slices in section 1. The selection process for original slices in section 2 follows a similar procedure, except that we change the $z_{glb}^{t_i}$ value with $z_{4,1}^{t_i}$. For original slices in section 3, there is a slight variation. When dealing with an original slice $z_{3,k}^{t_i}$ at $t_i$, since the section one section away from it is section 1, we consider z-slices below the minimum z-coordinate of section 2 as candidate negative samples. Consequently, we computed the $z_{lub}^{t_i}$ instead of the $z_{glb}^{t_i}$. The value of $z_{lub}^{t_i}$ is set to $z_{2,1}^{t_i}$, thus $z_{3,k}^{t_i}$ only considers z-slices between $z_L^{t_i}$ and $z_{lub}^{t_i}$ at $t_i$ as candidate negative samples As a result, case 1 remains unchanged, while case 2 uses linear interpolation between $(t_i, z_{2,1}^{t_i})$ and $(t_i + t_N, z_{3,k}^{t_i})$. The selection process for an original slice $z_{4,k}^{t_i}$ at $t_i$ follows a similar approach, except that we replace the $z_{lub}^{t_i}$ value with $z_{3,1}^{t_i}$. Moreover, original slices in section 3 and 4 at $t_i$ will not consider z-slices at $t_i$ for negative samples because those z-slices overlap with the negative samples produced by original slices in sections 1 and 2 at $t_i$.

***4. Downsampling and balancing negative samples.*** To enrich for informative negative samples rather than trivially dissimilar samples, we downsampled negative samples generated from case 1. We retained all samples generated by case 2, because these samples showed noticeable changes from the original slice without being overly different in overall shape. To balance the number of negative samples generated by case 1 and case 2, we applied a ratio-based downsampling procedure. If the number of negative samples produced by case 1 was at least twice the number of negative samples produced by case 2, we retained all negative samples generated by case 2 and randomly sampled twice the number of case 2 samples from the negative samples generated by case 1. If the number of negative samples produced by case 1 was less than twice the number produced by case 2, we retained both sets of samples.

Despite this reduction, the number of negative samples still exceeded the number of positive samples by a substantial margin. To further balance the positive and negative sample sets, we assigned a probability of 0.1 to each original slice when generating negative samples. This procedure randomly retained 10% of the negative samples generated through the procedure described above. Further reduction of sample redundancy is described in the section ***Further reducing sample redundancy***.

**Sensitivity analysis for temporal sample-selection parameterization.** To account for differences in developmental tempo across embryogenesis, STERN training used a time-dependent sampling parameterization in which positive and negative temporal intervals were specified separately for earlier and later developmental windows (Supplementary Table S2). In particular, the temporal interval parameters $t_A$ and $t_N$, defined above for positive and negative sample selection, were implemented separately for the first and second developmental windows of each embryo. These parameters are listed explicitly in Supplementary Table S2 using the notation $t_{A,1}$, $t_{A,2}$, $t_{N,1}$ and $t_{N,2}$, where subscripts 1 and 2 refer to the first and second developmental windows, respectively. The quantities $t_{shrink}$ and $z_{shrink}$ correspond to the temporal and spatial shrink factors described above. Because this time-dependent parameterization might influence sample selection between earlier and later developmental windows, we additionally trained a control model using a time-invariant sampling parameterization, where $t_{A,1}$ = $t_{A,2}$ and $t_{N,1} = t_{N,2}$ and these matched values are denoted as $t_A$ and $t_N$ in Supplementary Table S3. The resulting Embryo A similarity matrix remained highly consistent with that obtained using the time-dependent parameterization (Supplementary Fig. 7), including preservation of the broader high-similarity region observed during early developmental stages. These results indicate that the extended developmental continuity observed during early embryogenesis is not an artefact of the time-dependent sampling parameterization, but reflects intrinsic developmental organization captured by the learned representation.

**Further reducing sample redundancy**

For a given original slice $z_{i,k}^{t_i}$ in section *i* at $t_i$, nearby z-slices often exhibit minimal change in shape. As a result, the positive and negative samples generated from $z_{i,k}^{t_i}$ are frequently similar to those generated from neighbouring slices. This leads to redundant positive and negative samples. To address the issue of generating repetitive samples, we introduced a z-gap parameter when selecting the original slices, ensuring that two consecutive slices differ by the z-gap in their z-coordinates. This reduces the likelihood of selecting slices that are too similar in shape, thus minimizing redundant positive and negative samples. Determining an appropriate z gap automatically is difficult. To tackle this, we adopted a similar method to the one used for determining lower and upper bounds. Let $z_{gap,i}(t)$ be the z-gap for section *i* at the time $t$. We manually determined the value of $z_{gap,i}(t)$ at the start and final time points by inspecting the number of z-coordinate incremenets required for an image to show a noticeable change in shape. For intermediate time points, we used linear interpolation to estimate the z-gap. To further avoid redundant sampling, we determined the minimum number of original slices required for each section *i*, taking into account the computed z-gaps. Let the z-coordinate range of section *i* at time point *t* be defined by $s_{max,i}(t)$ and $s_{min,i}(t)$, which represent the maximum and minimum z-slices within section *i* at time *t*, respectively. For each time point *t*, we compute the following quantity for section *i*:

$$C_i(t) = \frac{s_{max,i}(t) - s_{min,i}(t)}{z_{gap,i}(t)}$$

We calculated $C_i(t)$ for every time point. The final value for section *i*, denoted by $N_{min,i}$, is the minimum of these computed values across all time points:

$$N_{min,i} = \min_t \left(C_i(t)\right)$$

To minimize redundancy, the next available original slice for an original slice at $z_{i,k}^{t_i}$ has a z-coordinate of $z_{i,k}^{t_i} + N_{min,i} \cdot z_{gap,i}(t_i)$ . By applying this procedure, we substantially reduced the number of redundant samples, enhancing the diversity of the dataset and ensuring that the selected slices better represent the variability within each section.

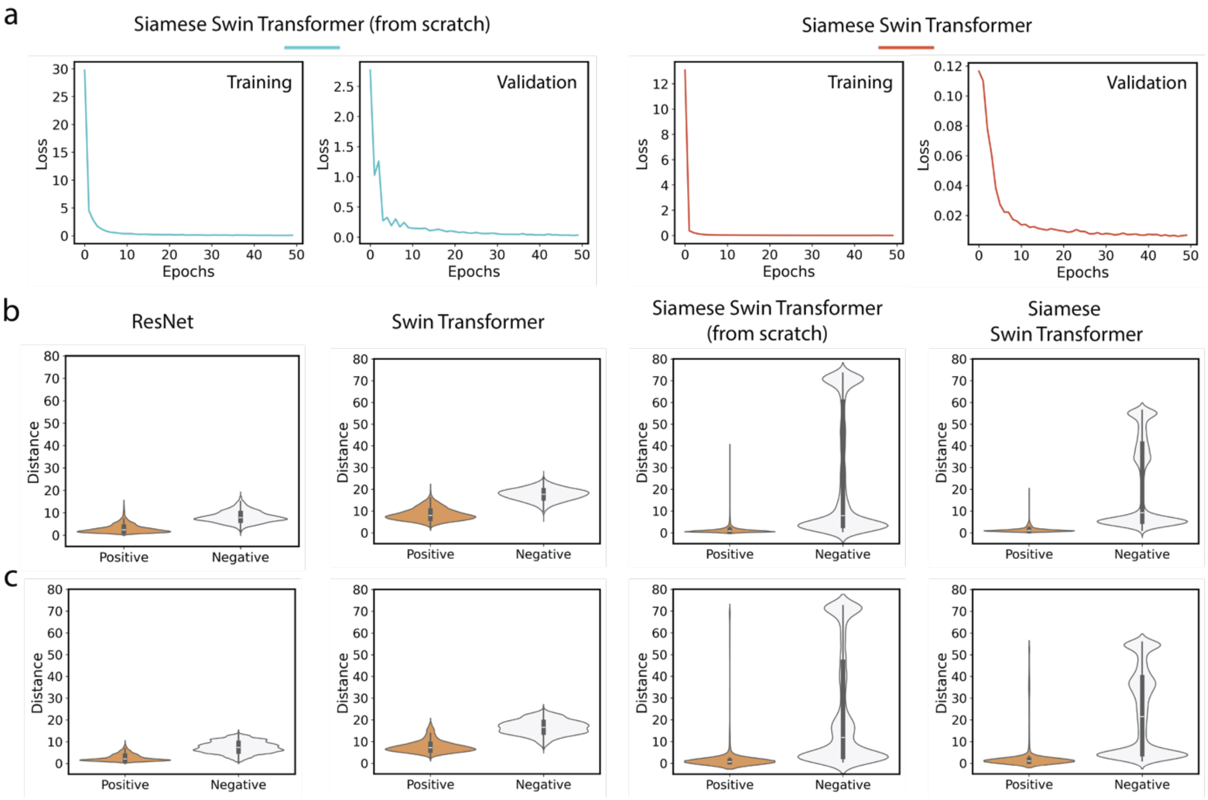


**Supplementary Fig. 1 | Training details and comparison of distance distributions in the test data. a,** Training loss and validation loss curves over 50 epochs for two variants of the Siamese Swin Transformer. The left panel shows the model trained from scratch with random initialization, while the right panel column shows the model initialized with pretrained weights. Both models converge, though the pretrained model (right) demonstrates lower initial loss values and smoother convergence compared to the from-scratch model (left), which exhibits higher initial loss values and more pronounced fluctuations during training. **b,** Distance distributions between positive (orange) and negative (gray) image pairs for four different model configurations on test-ABC dataset: ResNet (leftmost), Swin Transformer (center-left), Siamese Swin Transformer from scratch (center-right), and Siamese Swin Transformer (rightmost). Violin plots show the full distribution of distances, with internal box plots indicating median and quartile values. **c,** Corresponding distance distributions for the same models evaluated on test-D dataset. All violin plots within each panel share the same y-axis range for cross-model comparison.

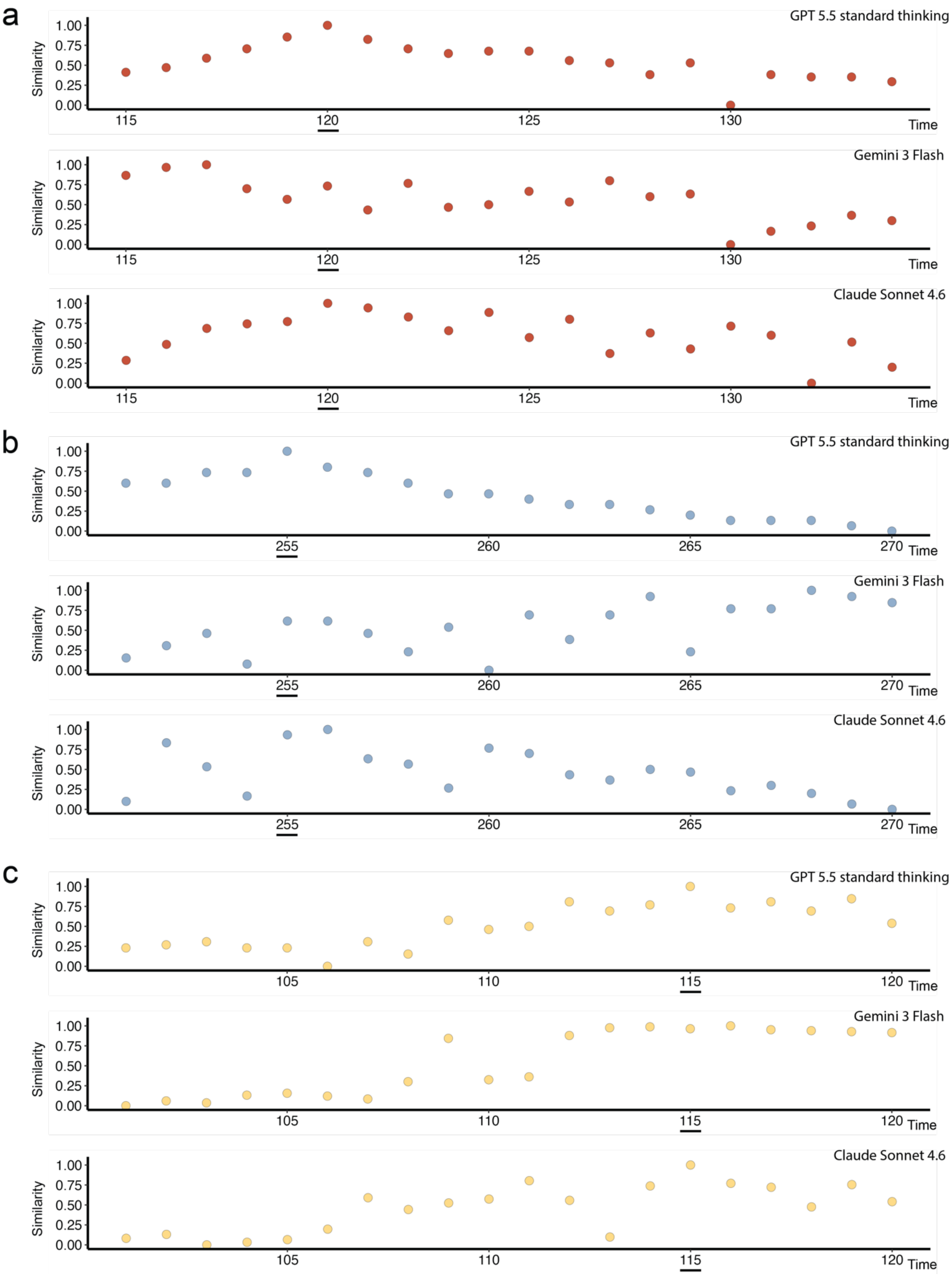


**Supplementary Fig. 2 | Individual multimodal AI model similarity trajectories.** Normalized similarity scores assigned by individual multimodal AI models across time for three heart datasets. **a**, Heart A; **b**, Heart B; **c**, Heart C. Each row corresponds to a single model (GPT-5.5 standard thinking, Claude Sonnet 4.6 or Gemini 3 Flash), and each point represents the similarity score assigned to one reference image. Scores were normalized independently within each model and dataset. The underlined time point indicates the query image.

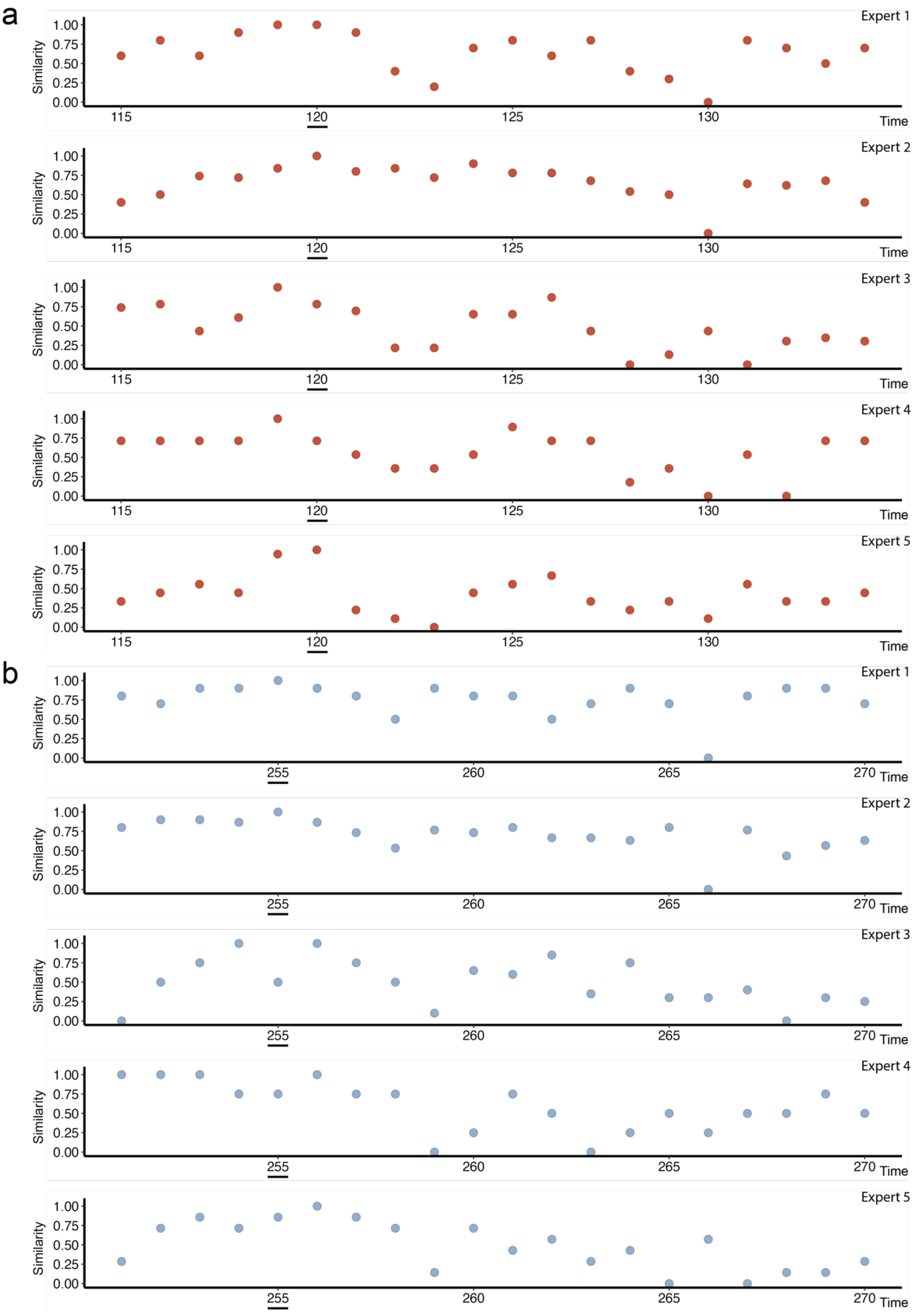

a
b
Expert 1
Expert 2
Expert 3
Expert 4
Expert 5
Similarity
Time
1.00
0.75
0.50
0.25
0.00
115
120
125
130
255
260
265
270

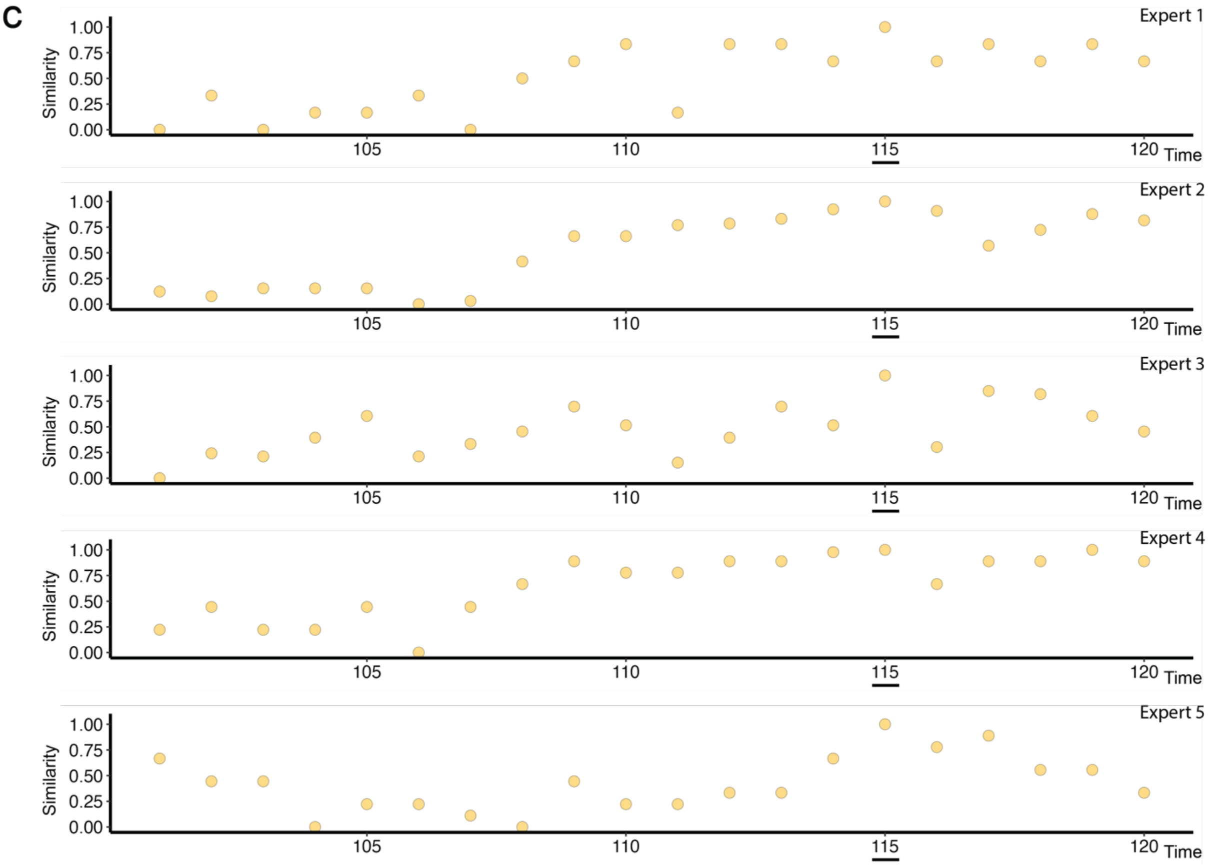


**Supplementary Fig. 3 | Individual expert similarity trajectories.** Normalized similarity scores assigned by individual domain experts across time for three heart datasets. **a**, Heart A; **b**, Heart B; **c**, Heart C. Each row corresponds to a single expert, and each point represents the similarity score assigned to one reference image. Scores were normalized independently within each expert and dataset. The underlined time point indicates the query image.

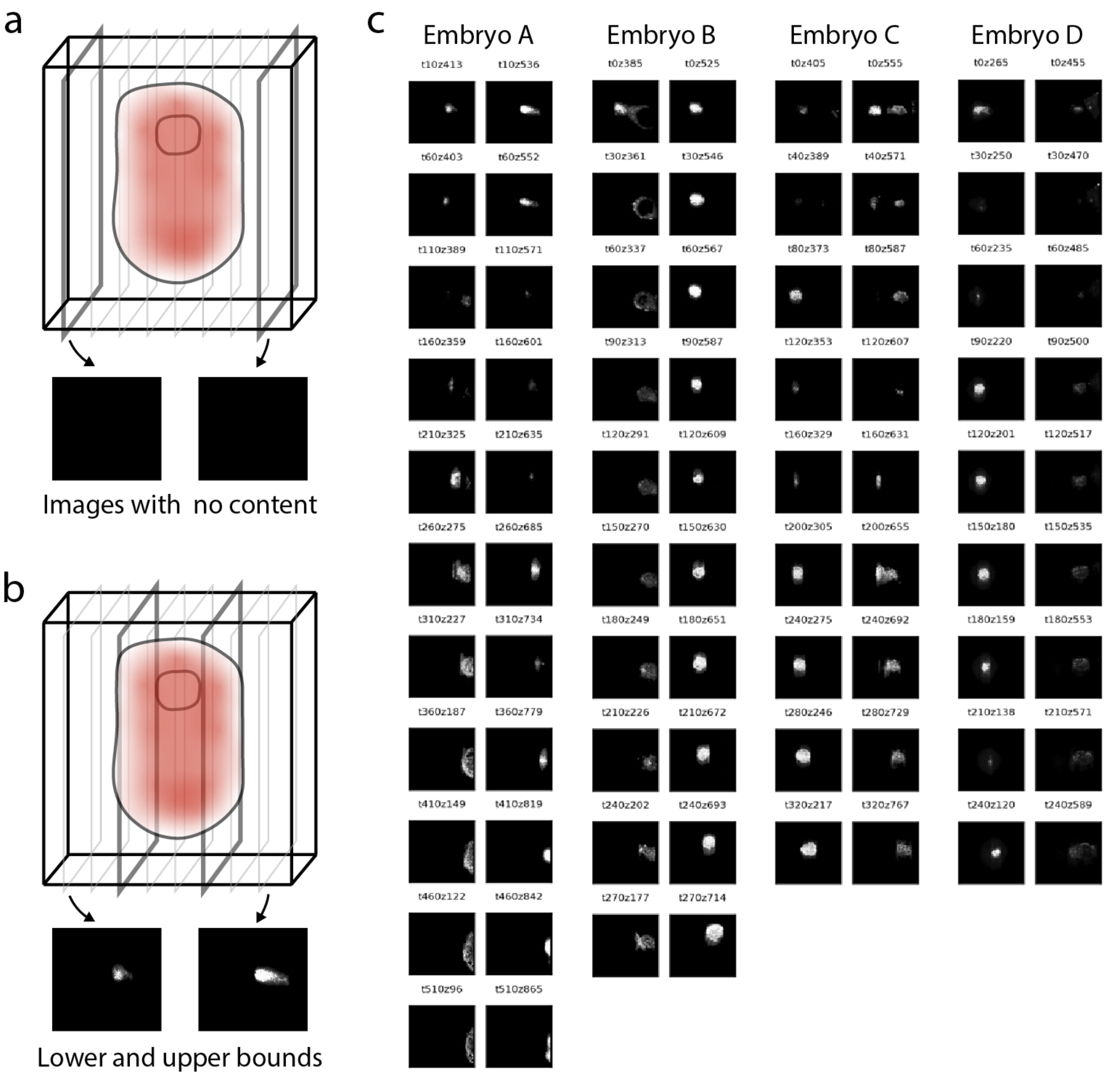


**Supplementary Fig. 4 | Methodology for determining and validating z-slice bounds in embryonic imaging data. a,** Schematic illustration showing how empty image slices (shown in black) occur outside the embryonic region in the 3D imaging volume. **b,** Illustration demonstrating the concept of upper and lower z-slice bounds with example images. **c,** Validation of the bound selection approach across four different embryos (A-D). For each embryo, pairs of images show the lower and upper bounds (left and right respectively) at selected time points that span the entire developmental period. Each image is labelled with its temporal (t) and z-coordinate information.

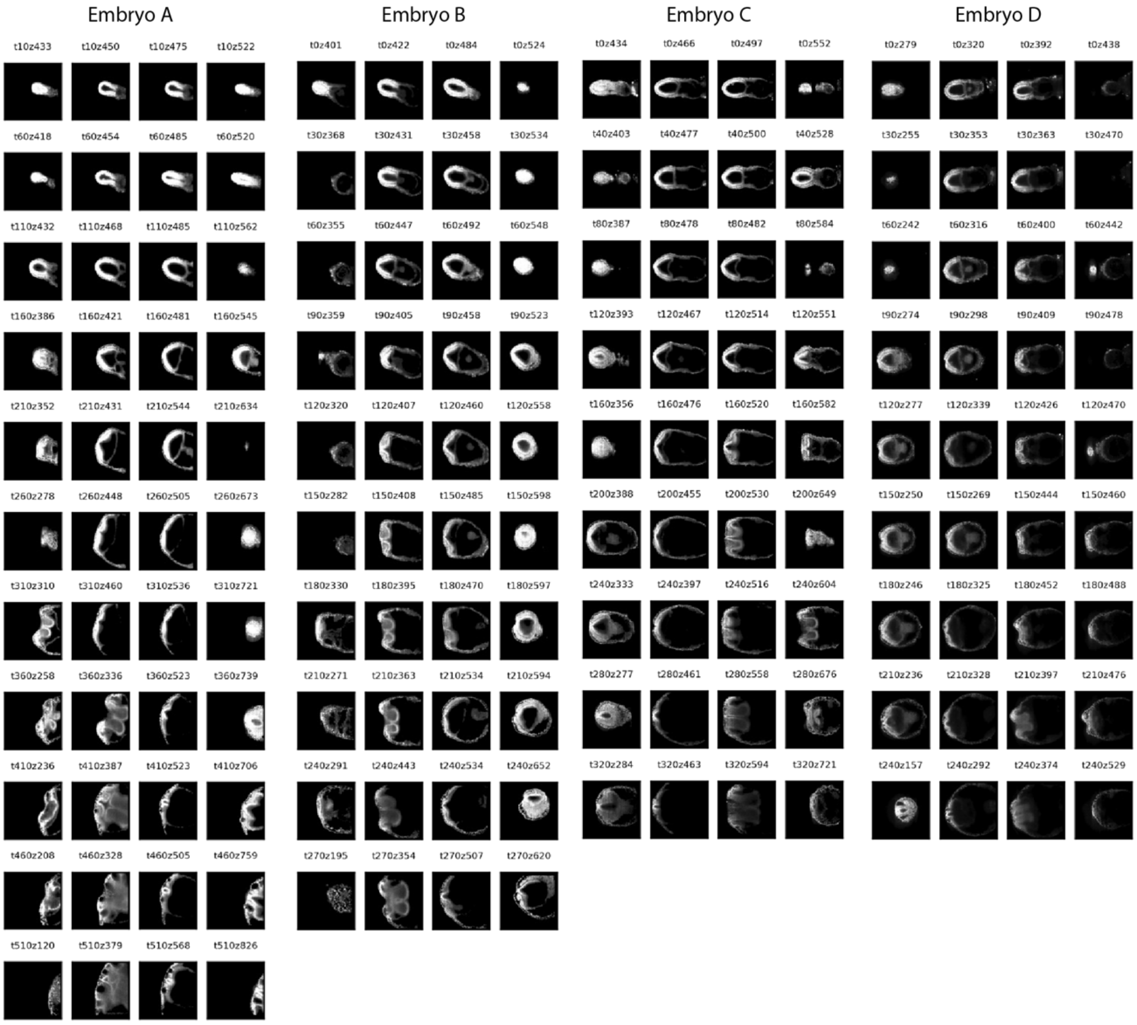


**Supplementary Fig. 5 | Visualization of embryo content across different z-slice sections.** Representative images from four embryos (A-D) showing the division of z-slices into four sections with equal numbers of slices. For each embryo at different time points, four images are shown where, from left to right: the first image is randomly selected from section 1, the second from section 2, the third from section 3, and the fourth from section 4. Each image is labelled with its temporal (t) and z-coordinate information.

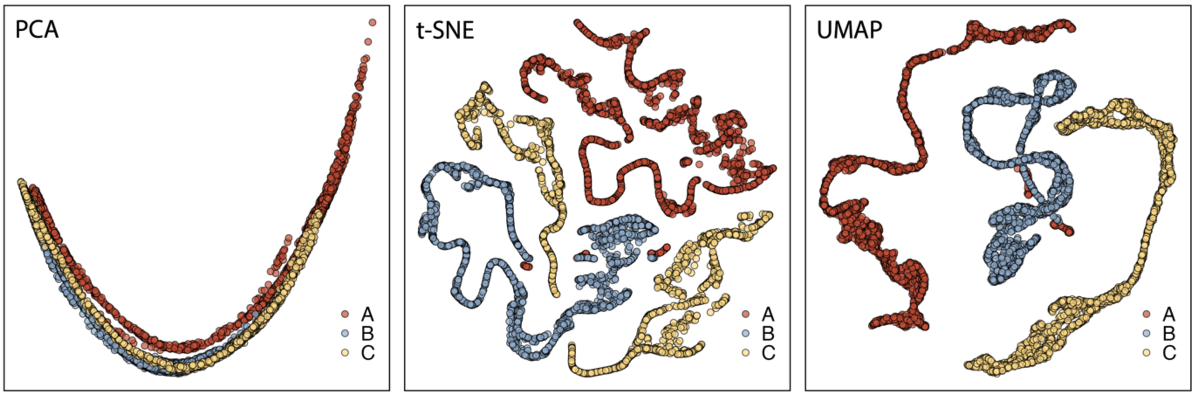


**Supplementary Fig. 6 | Dimensionality reduction analysis of three heart datasets.** Visualization using three reduction methods on Hearts A (red), B (blue), and C (yellow). (Left) PCA projection to 2D. (Middle) t-SNE with a perplexity of 40 and 300 iterations. (Right) UMAP using '*n_neighbors=40*' and '*min_dist=0.6*'.

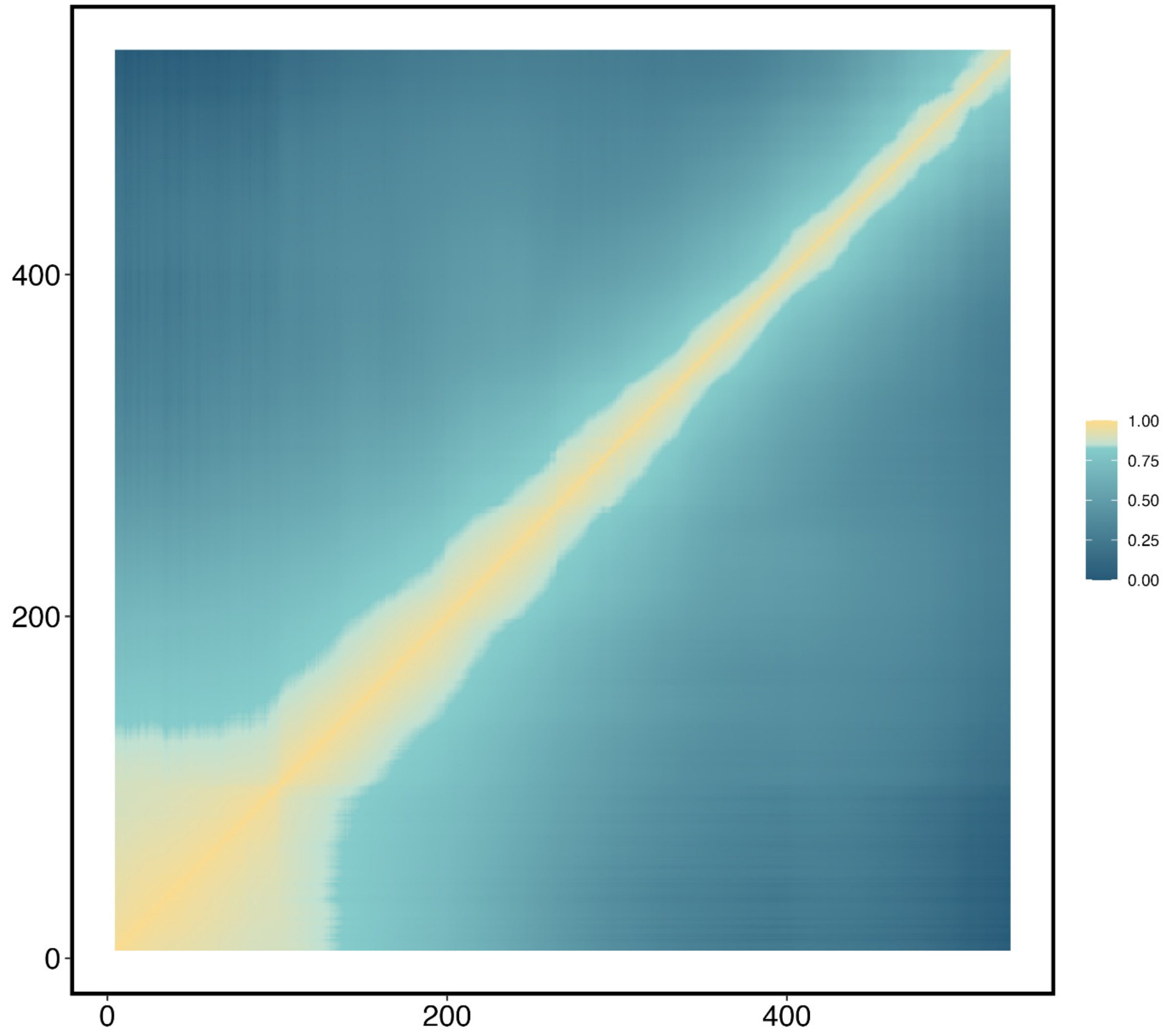


**Supplementary Fig. 7 | Embryo A similarity matrix from sensitivity analysis using a time-invariant sampling parameterization.** Embryo A similarity matrix obtained from the control model using the time-invariant sampling parameterization, in which first- and second-window temporal interval values were matched within each embryo, as listed in Supplementary Table S3.

| **Expert ID** | Area of expertise | Years of experience |
| --- | --- | --- |
| Expert 1 | Mouse genetics and developmental biology, focused on growth regulation | 35 years |
| Expert 2 | Bioimage analysis and microscopy; cell biology, specifically cell migration and morphology | 25 years |
| Expert 3 | Intravascular imaging, intravascular ultrasound (IVUS), optical coherence tomography (OCT) | 3 years |
| Expert 4 | Spatial transcriptomics data analysis, computational biology, single-cell and spatial multi-omics modelling | 3 years |
| Expert 5 | Clinical cardiac electrophysiology and arrhythmia research | 5 years |

**Supplementary Table S1 | Background information for domain experts participating in the blinded similarity evaluation.** Areas of expertise and years of experience are reported as provided by each participant. Expert identities are anonymized.

| **Embryo** | $t_{start,1}$ | $t_{A,1}$ | $t_{start,2}$ | $t_{A,2}$ | $t_{shrink}$ | $z_{shrink}$ | $t_{N,1}$ | $t_{N,2}$ |
|---|---|---|---|---|---|---|---|---|
| A | 5 | 20 | 263 | 25 | 2 | 4 | 50 | 100 |
| B | 0 | 20 | 143 | 25 | 2 | 3 | 50 | 50 |
| C | 0 | 20 | 175 | 25 | 2 | 4 | 50 | 100 |
| D | 0 | 20 | 133 | 25 | 2 | 3 | 50 | 40 |

**Supplementary Table S2 | Time-dependent sampling parameterization used for STERN training.** Temporal interval parameters were specified separately for the first and second developmental windows of each embryo. $t_{start,1}$ and $t_{start,2}$. define the starting time points of the two windows. For each original slice, the temporal parameters were selected according to the developmental window containing its time point $t_i$. $t_{A,1}$ and $t_{A,2}$ denote the first- and second-window values of $t_A$ for positive sample selection, and $t_{N,1}$ and $t_{N,2}$ denote the corresponding values of $t_N$for negative sample selection. The quantities $t_{shrink}$ and $z_{shrink}$ correspond to the temporal and spatial shrink factors described in the Supplementary Methods.

| **Embryo** | $z_{shrink}$ | $t_A$ | $t_{shrink}$ | $t_N$ |
|---|---|---|---|---|
| A | 4 | 20 | 2 | 50 |
| B | 3 | 20 | 2 | 50 |
| C | 4 | 20 | 2 | 50 |
| D | 3 | 20 | 2 | 40 |

**Supplementary Table S3 | Time-invariant sampling parameterization used for sensitivity analysis.** This control parameterization used matched first- and second-window temporal interval values within each embryo to assess whether the observed developmental structure depended on the time-dependent sampling parameterization used for STERN training. In this setting, $t_{A,1}$ = $t_{A,2}$ and $t_{N,1} = t_{N,2}$ so the matched values are denoted as $t_A$ and $t_N$.

**Supplementary Note 1 | Multimodal AI similarity evaluation prompt**

For the blinded multimodal AI evaluation, each model was given one query image and 20 randomized reference images from the same heart dataset. Images were uploaded in three batches, and models were instructed not to score images until all files had been uploaded.

The evaluation was performed using three contemporary general-purpose multimodal AI systems accessed through their public web interfaces: GPT-5.5 standard thinking via ChatGPT (https://chatgpt.com/), Claude Sonnet 4.6 via Claude (https://claude.ai/) and Gemini 3 Flash via Gemini (https://gemini.google.com/). Model names were recorded as displayed in the corresponding interfaces at the time of evaluation.

The exact prompt structure used for all multimodal AI evaluations is provided below.

Phase 1 — Rules & protocol
"
*Role: You are an expert in developmental biology and morphological assessment.*

*As part of an evaluation focused on morphological similarity in high-resolution images of developing mouse hearts, you will assess subtle structural differences that evolve continuously over short developmental time scales.*

*You will be given:*
*1. One query image.*
*2. Twenty reference images from one independent evaluation set.*

*The images will be uploaded in three batches:*
*Batch 1: the query image.*
*Batch 2: the first 10 reference images.*
*Batch 3: the remaining 10 reference images.*

*Please do not analyse or score any images until all files have been uploaded.*

*After all files have been uploaded, please enter a similarity score from 1 to 100 for each image.*

*Please do not rename image files, as filenames are used to match scores automatically.*

*Output format:*
*Return only a CSV-formatted code block with exactly these columns:*

*image_name,similarity_score*
"

Phase 2 — upload prompts
"
*Here is Batch 1: the query image.*
*Please hold this in memory. Do not analyse yet.*

*"*

*"*
*Here is Batch 2: the first 10 reference images.*
*Please hold these in memory. Do not analyse yet.*
*"*

*"*
*Here is Batch 3: the remaining 10 reference images.*
*Please hold these in memory. Do not analyse yet.*
*"*

Phase 3 — trigger prompt
*"*
*ALL FILES UPLOADED.*

*Please now enter your similarity scores (1–100) in the similarity_score column corresponding to each image name.*

*Return only a CSV-formatted code block with exactly these columns:*

*image_name,similarity_score*
*"*
